# Data-driven analysis of COVID-19 reveals specific severity patterns distinct from the temporal immune response

**DOI:** 10.1101/2021.02.10.430668

**Authors:** Jackwee Lim, Kia Joo Puan, Liang Wei Wang, Karen Wei Weng Teng, Chiew Yee Loh, Kim Peng Tan, Guillaume Carissimo, Yi-Hao Chan, Chek Meng Poh, Cheryl Yi-Pin Lee, Siew-Wai Fong, Nicholas Kim-Wah Yeo, Rhonda Sin-Ling Chee, Siti Naqiah Amrun, Zi Wei Chang, Matthew Zirui Tay, Anthony Torres-Ruesta, Norman Leo Fernandez, Wilson How, Anand K. Andiappan, Wendy Lee, Kaibo Duan, Seow-Yen Tan, Gabriel Yan, Shirin Kalimuddin, David Chien Lye, Yee-Sin Leo, Sean W. X. Ong, Barnaby E. Young, Laurent Renia, Lisa F.P. Ng, Bernett Lee, Olaf Rötzschke

**Author notes:** First authors (Jackwee Lim, Kia Joo Puan, Liang Wei Wang). Senior authors (Laurent Renia, Lisa F.P. Ng, Bernett Lee, Olaf Rötzschke). Correspondence (O.R.).

## Abstract

Key immune signatures of SARS-CoV-2 infection may associate with either adverse immune reactions (severity) or simply an ongoing anti-viral response (temporality); how immune signatures contribute to severe manifestations and/or temporal progression of disease and whether longer disease duration correlates with severity remain unknown. Patient blood was comprehensively immunophenotyped via mass cytometry and multiplex cytokine arrays, leading to the identification of 327 basic subsets that were further stratified into more than 5000 immunotypes and correlated with 28 plasma cytokines. Low-density neutrophil abundance was closely correlated with hepatocyte growth factor levels, which in turn correlated with disease severity. Deep analysis also revealed additional players, namely conventional type 2 dendritic cells, natural killer T cells, plasmablasts and CD16^+^ monocytes, that can influence COVID-19 severity independent of temporal progression. Herein, we provide interactive network analysis and data visualization tools to facilitate data mining and hypothesis generation for elucidating COVID-19 pathogenesis.

## Introduction

The pandemic coronavirus disease 2019 (COVID-19) is caused by the severe acute respiratory syndrome coronavirus 2 (SARS-CoV-2). As of 25^th^ January 2021, there are 100 million cases with more than 2 million deaths worldwide (Dong et al., 2020). While the immunogenic responses in COVID-19 vaccinated individuals remain largely unknown, the majority of the COVID-19 patients are asymptomatic or have mild flu-like symptoms, some develop severe diseases, which may lead to death from respiratory failure or even multi-organ failure from a hyperactivated immune system (Manson et al., 2020). The immunopathology in severe COVID-19 is characterized by lymphopenia, a sustained loss of CD4^+^ and CD8^+^ T cells, a dramatic increase in the number of immature neutrophils as well as an altered myeloid cell compartment in severe COVID-19 cases (Carissimo et al., 2020; Jouan et al., 2020; Kuri-Cervantes et al., 2020; Silvin et al., 2020). Another hallmark of severe COVID-19 is the cytokine storm associated with elevated levels of cytokines (IL1β, IL1Rα, IL2, IL7, IL10, G-CSF, TNFα), chemokines (IP10, MCP1, MIP1α) and endogenous neutrophil calprotectin (Ragab et al., 2020; Silvin et al., 2020; Barnaby E. Young et al., 2020).

Although many studies have independently identified different immunotypes and cytokines associated with SARS-CoV-2 infection and severe COVID-19, their interplay is still largely unknown for distinguishing active infection, severity and post-infection aberrations. In this work, we analyzed the blood samples of 77 hospitalized patients and 10 healthy controls. Also, to obtain a more full-bodied approach, we have compiled the data and provided a comprehensive COVID-19 resource, which integrates high dimensional mass cytometry and bead-based cytokine/chemokine arrays. Network analyses between the key immune subsets and the associated cytokine levels in the plasma were used to correlate clinical states and disease trajectory of COVID-19. One of the key findings is the identification of a link between hepatocyte growth factor (HGF) and low-density neutrophils (LD Neu). While both factors are closely associated with disease severity, it represents only the tip of the iceberg; the network of underlying pathways is complex and involves many players. By sharing these data together with easy-to-use visualization tools, we provide a rich resource to study the manifestation of COVID-19 as well as the particulars of the SARS-CoV-2 induced immune response.

## Results

### Study design and clinical characteristics of the cohort

To determine the changes induced by SARS-CoV-2 infection in peripheral blood mononuclear cell (PBMC) subsets, we assembled a comprehensive mass cytometry panel of 40 antibodies against lineage-specific markers, adhesion molecules and other surface molecules indicative of the functional state of the cells (Figure 1A). This panel divided the entire PBMC population into 327 partly overlapping subsets, which consisted of both extensively described immune cell populations such as T cells, B cells, dendritic cells, monocytes, natural killer (NK) cells, basophils, mucosal-associated invariant T (MAIT) cells, innate lymphoid cells (ILCs) and LD Neu, as well as lesser-known immune cell populations such as CD56^+^ monocytes, CD56^+/-^ MAIT and the PD-L1^+^ LD Neu subsets (Villani et al., 2017). Cytokine bead arrays based on Luminex™ and ultra-sensitive Quanterix™ technologies were used to quantify the changes of twenty-eight cytokines in patient plasma (Figure 1A). The interconnected databases generated by these approaches were then subjected to network analysis to determine immune signatures of the anti-SARS-CoV-2 response as well as the cellular and molecular drivers of severe COVID-19.

**Figure 1.**
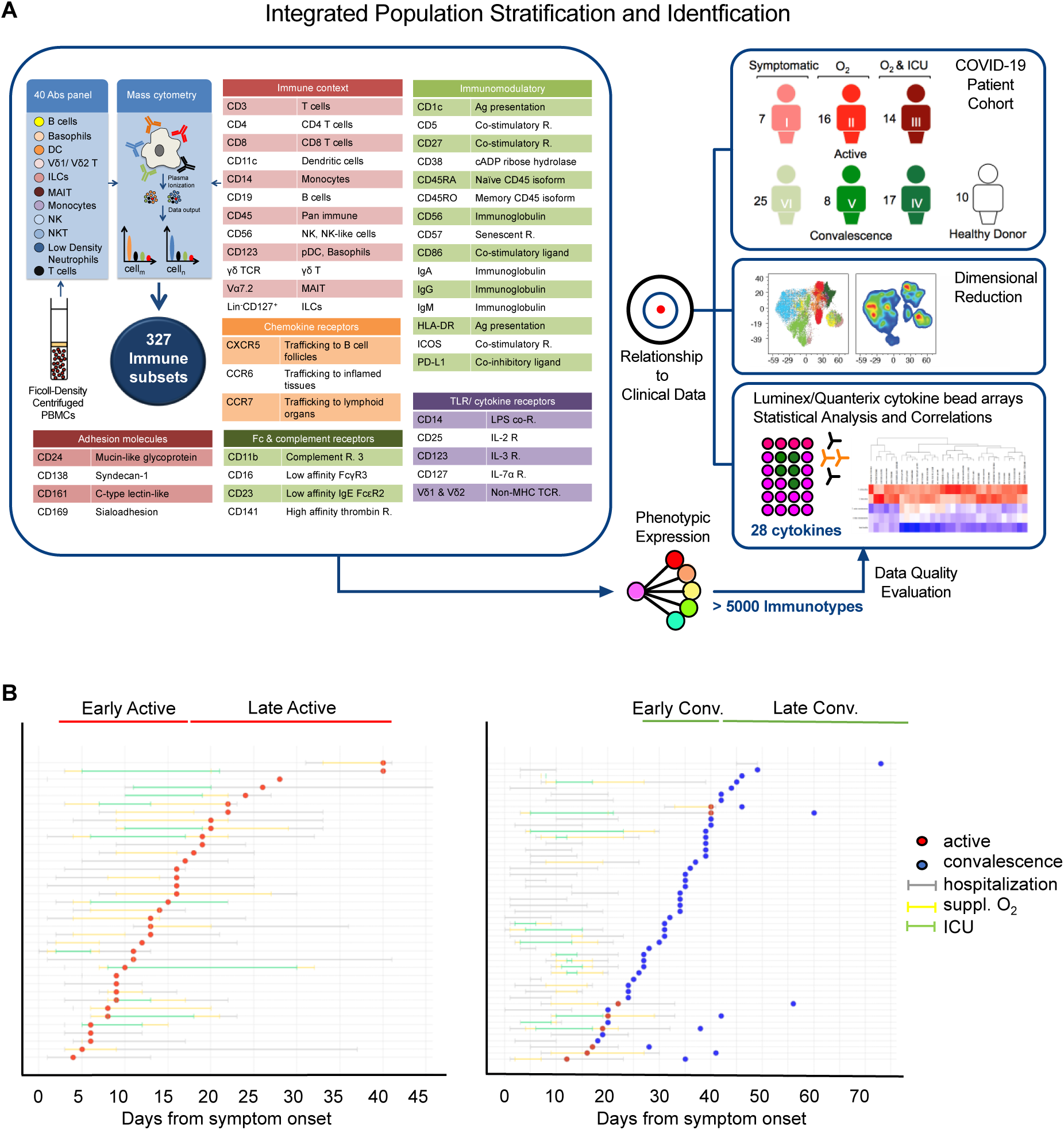
Study design and clinical characteristics of the cohort. (A) Schematic showing the pipeline for sample acquisition and analysis. A list of the antibody targets is presented. (B) Timelines for individual COVID-19 cases, indicating points of sample collection and any clinically pertinent detail e.g. duration of hospitalization, oxygen supplementation and admission to the intensive care unit (ICU).

In order to establish a timeline on the SARS-CoV-2 specific immune response, 87 blood samples were collected from 77 hospitalized COVID-19 patients at varying time points (Figure 1B). The state of infection was determined by SARS-CoV-2 real-time reverse transcriptase polymerase chain reaction (PCR) as previously described (Corman et al., 2020; Barnaby E Young et al., 2020). PCR positive samples were grouped into early active (median days post illness onset (PIO): 9) and late active (median days PIO: 20). PCR negative samples were grouped into early convalescence (median days PIO: 25) and late convalescence (median days PIO: 39) (Figure 1B). As a proxy for disease severity, the samples were stratified based on treatment regime into three severity groups, namely mild (symptomatic; n = 32), moderate (symptomatic with supplemental oxygen (suppl. O_2_); n = 24) and severe (suppl. O_2_ and need for intensive care unit (ICU); n = 31). With the exception of age and gender, most demographic variables did not differ significantly between COVID-19 patients and healthy donors, as well as between severity groups (Table S1). Severity was associated with advanced age (p < 0.0001).

### Temporal variations in PBMC subsets during active and convalescent COVID-19

Mass cytometry analysis allowed gating of total PBMCs into 38 non-overlapping basic subsets from both the adaptive and innate arms of the immune system (Figure 2A and Table S2). Uniform Manifold Approximation and Projection (UMAP) analysis segregated these cells into distinct clusters of T cells, B cells, NK cells, monocytes and other antigen presenting cells (Figure 2A, left). Surprisingly, large numbers of LD Neu were detected (Figure 2A, right). Neutrophils are typically not present in the PBMC fraction of healthy donors but found in abundance in COVID-19 samples. Subsequent analysis with a targeted mass cytometry panel confirmed that these cells are indeed LD Neu by revealing an immunophenotype that is consistent with neutrophils (CD66b^+^CD15^+^CD16^high^CD10^+^CD24^+^) (Table S2).

**Figure 2.**
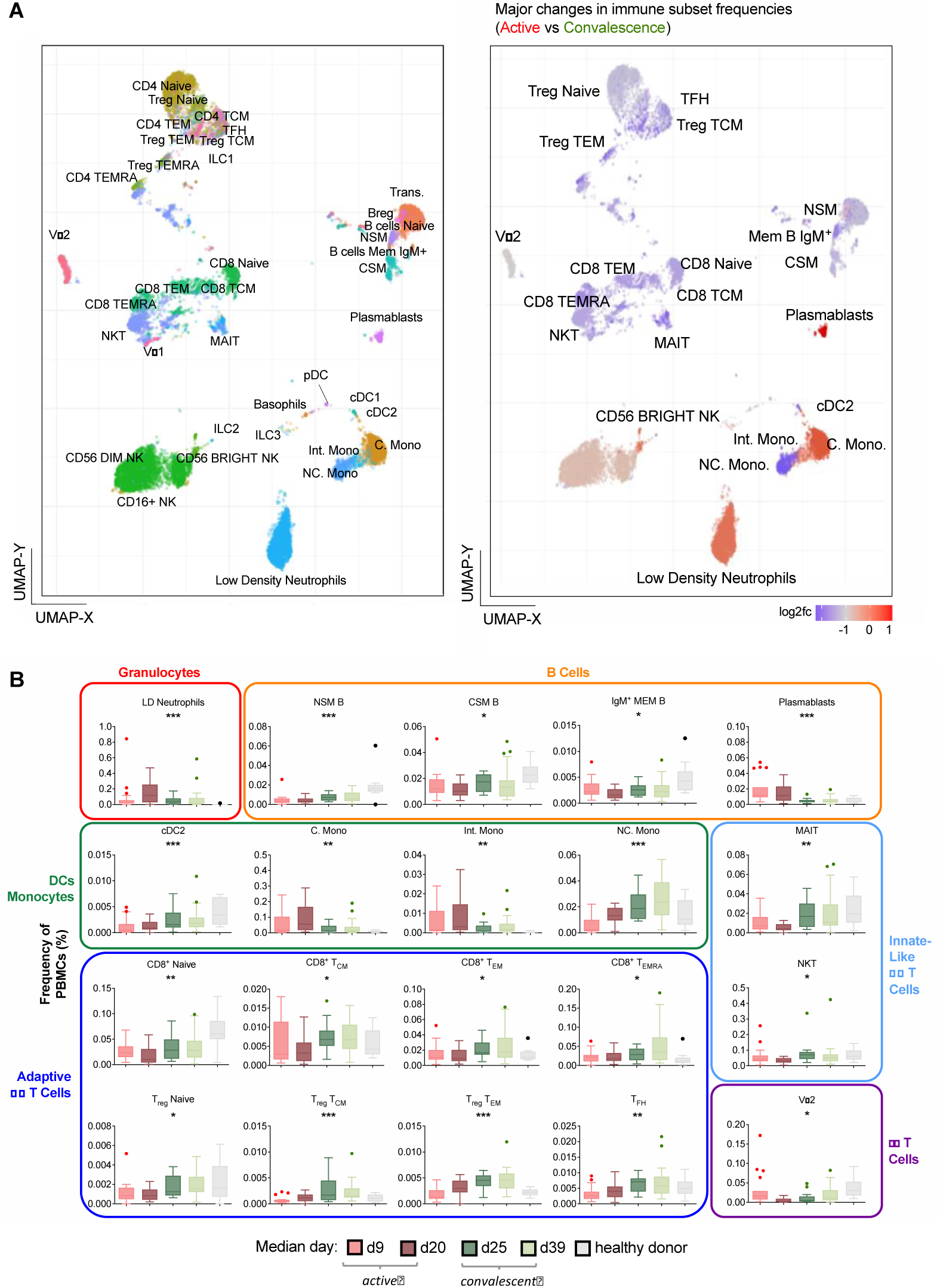
Frequency changes in 38 basic immune cell populations with SARS-CoV-2 infection. (A) Uniform Manifold Approximation and Projection (UMAP) plots of 38 main immune cell populations detected by mass cytometry (left). Right: heat map representation of fold changes in the detected populations during active infection relative to convalescence, with color-coding done on a log_2_ scale. (B) Frequency-time plots of immune cell populations of interest over the course of disease.

Generalized lymphopenia was observed in the active phase relative to convalescent COVID-19 (Figure 2A, right), particularly in various CD4^+^ and CD8^+^ T cell, most B cell, NKT and MAIT cell subsets. A significant reduction was also observed for non-classical monocytes (NC. Mono), while increased frequencies were detected for plasmablasts, classical monocytes (C. Mono) and LD Neu. Closer inspection of the temporal profiles for various subsets revealed a number of interesting trends (Figure 2B). For LD Neu, the increase was largely restricted to the late active phase, while the plasmablast frequency increased during the early active phase. The frequency of C. and intermediate (Int.) monocytes rose throughout active infection before declining in the convalescent phase. In contrast, the frequency of NC. Mono in the blood dropped sharply during the early active phase but increased gradually until late convalescence (Figure 2B).

The timeline of events is particularly evident in a heatmap display (Figure 3A, left). Based on the increase in subset frequencies, the earliest responders were NK cells followed by plasmablasts, C./Int. Mono and LD Neu. Further, there was also a sharp decrease in numbers, which may indicate the mobilization of subsets such as NC. Mono, T regulatory (Treg), T follicular helper (T_FH_) and CD4^+^ T cells to the lung mucosa as front-line responses. Their frequencies increased later during convalescence (Figures 2B and 3A). We also observed a loss in blood MAIT cell frequency during active infection that was restored to healthy levels during convalescence (Figure 3A), which is in line with a previous study suggesting active recruitment of MAIT to the lung mucosa (Parrot et al., 2020). Also the robust increase in the frequency of circulating plasmablasts during the active phase is consistent with previous single-cell analysis of patients with severe COVID-19 (Wilk et al., 2020). In contrast, non-class-switched memory (NSM), class-switched memory (CSM) and IgM^+^ memory B cells showed decreased frequencies, which increased during convalescence but did not recover to healthy levels.

**Figure 3.**
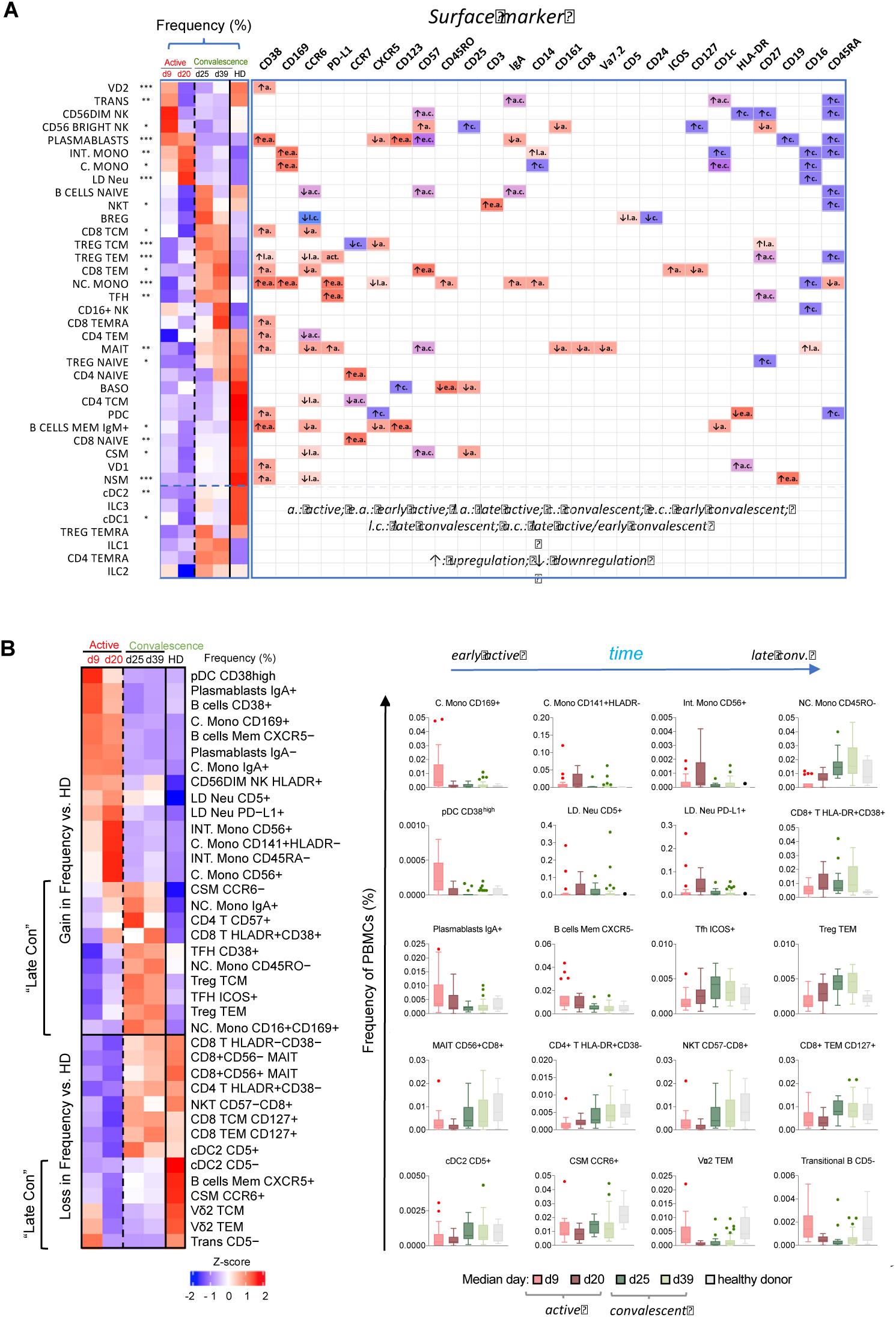
Temporal changes in frequencies and surface marker expression profiles of various immunotypes during active and convalescent COVID-19. (A) Left: heat map representation of the frequencies of all 38 basic immune cell populations as a function of days post illness onset (PIO) – d9 (early active; median: 9 days PIO), d20 (late active; median: 20 days PIO), d25 (early convalescence; median: 25 days PIO), d39 (late convalescence; median: 39 days PIO). Asterisks indicate statistical significance - *, p<0.05; **, p<0.01; ***, p<0.001 (one-way ANOVA of all disease phases and healthy controls). Right: up- or down-regulation of indicated surface markers for the 38 main immune cell populations as a function of disease phase. (B) Left: heat map representation of the frequencies of the top 38 immunotypes as a function of disease phase. Right: Box-and-whiskers plots of select immunotypes showing the frequency-time relationships, with mean and IQR indicated. “Late Con” refers to a group of immune subsets, which fail to recover to healthy levels even in late convalescence as post-infection aberrations.

Activation and differentiation of immune cells during the course of the SARS-CoV-2 infection were indicated by changes in the expression levels of surface markers (Figure 3A, right). The profiles of more than 5000 immunotypes derived from the 38 basic immune subsets can be assessed by an interactive online viewer (see Materials and Methods: Data and Code Availability and Table S2). Although expression of functional markers was highly heterogeneous across immune subsets and disease phases, we observed some notable trends. CD38 was upregulated across diverse immune subsets during active infection, indicative of widespread cell activation in response to viral infection. CD169, an adhesin that binds sialic acid, was also selectively upregulated in all monocyte subsets during the early but not late active phase. Many T and B cell subsets – Treg TEM, NSM and CSM cells – downregulated the lymph node homing molecule CCR6 over the course of disease, primarily during active infection, which may impede homing to inflamed tissues and development of germinal center responses (Figure S1). A range of cell subsets exhibited elevated levels of CD57 from early active infection to early convalescence, indicative of immune senescence (Figure 3A). Finally, CD16 (FcγRIII) was upregulated on MAIT cells in late active infection and on LD Neu, all monocyte subsets and CD16^+^ NK cells during convalescence (Figure 3A).

Based on the observed phenotype variations, we defined and curated 327 partly overlapping immunotypes and compared the frequencies of these subsets in each phase of COVID-19 (Figure 3B and Table S3). High levels of CD169^+^ C. Mono were observed exclusively during early active infection. The same also applied for CD38^high^ plasmacytoid dendritic cell (pDC) subsets and IgA^+^ plasmablasts, followed by IgA^-^ plasmablasts. Notably, four subsets of monocytes, namely CD141^+^HLADR^-^ C. Mono, CD56^+^ C. Mono, CD45RA^-^ Int. Mono and CD56^+^ Int. Mono peaked during late active SARS-CoV-2 infection and may portend recovery (Figure 3B). The increase in the LD Neu population from early to late active infection is reflected in both its PD-L1^+^ and CD5^+^ subsets. The active-phase lymphopenia affected virtually all CD8^+^ central (CD8 TCM) and effector memory T subsets (CD8 TEM) (Figure 3B). Significant losses of CD56^+^CD8^+^ MAIT, CCR6^+^ CSM, HLA-DR^+^CD38^-^ CD4^+^ T, CD57^-^CD8^+^ and NKT subsets were also observed during active infection (Figure 3B). Next, in convalescence, a frequency increase persisting into late phase was observed for T cells (such as CD57^+^CD4^+^ T, HLA-DR^+^CD38^+^ CD8^+^ T, CD38^+^/ICOS^+^ T_FH_ defined as circulatory CXCR5^+^, Treg TCM/TEM), NC. Mono (IgA^+^, CD45RO^-^ and CD16^+^CD169^+^) and CCR6^-^ CSM above healthy levels. Also in this phase, the loss of CD5^-^ conventional type 2 DC (cDC2), Vδ2 TCM, Vδ2 TEM, B cells memory CXCR5^+^, CCR6^+^ CSM and CD5^-^ transitional B cells persisted into late convalescence below healthy levels (Figure 3B). These late-convalescent immunotypes, which do not recover to healthy levels, may represent post-infection aberrations.

### Alterations of immunotypes associated with disease severity

Depending on the treatment given, active and convalescent patients were stratified into ‘mild’ (symptomatic), ‘moderate’ (symptomatic with suppl. O_2_) and ‘severe’ (suppl. O_2_ and ICU admission). Thus we obtained groups I, II and III denoting active mild, active moderate, and active severe, respectively and groups IV (severe), V (moderate), and VI (mild) for patients in the convalescent phase (Figure 1A). The global changes in the cell subset composition are particularly evident in UMAP plots colored according to the fold change in reference to healthy controls (Figure 3A). The most striking association was observed for LD Neu. While a minor increase was observed in group I (active, mild), the numbers increased dramatically in group II (active, moderate) and peaked in group III (active, severe). The numbers persisted in group IV (conv., severe) and gradually declined from group V (conv., moderate) to group VI (conv., mild) (Figures 4A, 4B and S2A). This was also reflected in the increased LD Neu-to-lymphocyte ratios compared with group VI mild convalescent and healthy (Figure S3A). The expansion of LD Neu was due to CD16^+^ neutrophils and CD16^high^ neutrophils. CD16^high^ neutrophils (Figures S3B and S3C) described herein resemble pseudo-Pelger-Huet cells, which were previously reported in other severe viral infections (Morrissey et al., 2020).

**Figure 4.**
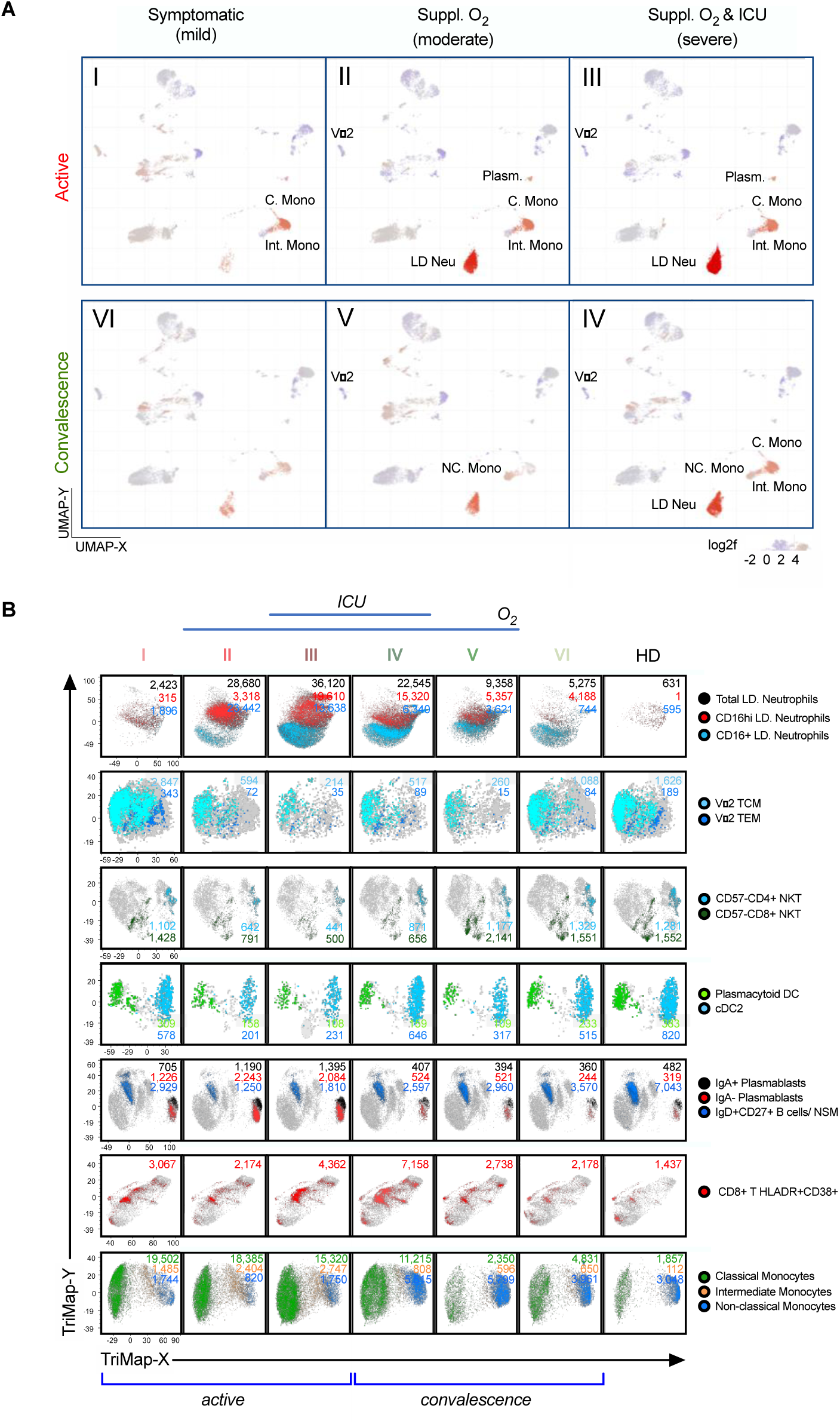
Alterations of immunotypes associated with the six-group disease severity states. (A) Distribution of 38 immune cell subsets among group I active mild symptomatic, group II active suppl. O_2_ group III active suppl. O_2_ ICU, group IV convalescent suppl. O_2_ ICU, group V convalescent Suppl. O_2_ and group VI convalescent mild symptomatic using UMAP clustering. Color indicates the log2 fold change in the frequency against healthy donors. (B) TriMap clustering of PD-L1^+^ LD Neu, Vδ2 TCM, Vδ2 TEM, pan-CD57^-^ NKT, pDC, cDC2, IgA^+^/^-^ plasmablasts, IgD^+^CD27^+^ NSM, HLA-DR^+^CD38^+^ CD8 T cells, C. Mono, Int. Mono and NC. Mono among 6 groups of SAR-CoV-2 patients and healthy donors (HD). The absolute number shown for the immune cells has been normalized per 300,000 PBMCs and thus reflects its frequency.

Two other immune cell subsets, which closely associated with severe COVID-19 are Vδ2 T and NKT cells (Figures 4B and S2A). Their frequencies sharply declined in group II/III/IV patients. Further analysis revealed that the loss of Vδ2 T is likely due to Vδ2 TCM and Vδ2 TEM, which remained significantly reduced in frequency in groups IV/V compared with healthy (Figures 4B, 5A and S4A). Similarly for NKT sub-populations, the CD57^-^ NKT cell but not CD57^+^ NKT cell subset was reduced in frequency in groups II/III/IV (Figure S4D). The loss of the CD57^-^ NKT cells included the CD4^+^, CD8^+^ and the double-negative (DN) NKT sub-populations. Although reduced numbers were also detected for dendritic cell subsets, the frequencies of pDC and cDC2 were only marginally suppressed across groups II/III/IV/V (Figures 4B, S4B and S4C). For plasmablasts, a subtle increase in frequency was observed in the active phase groups I to III but waned during convalescence (Figures 4A, 4B, S5A, and S5B). Notably, IgA^-^ plasmablasts showed a higher increase compared with its IgA^+^ counterpart (6.53-fold vs 2.89-fold compared with HD) in group III (Figures 4B and S5B).

**Figure 5.**
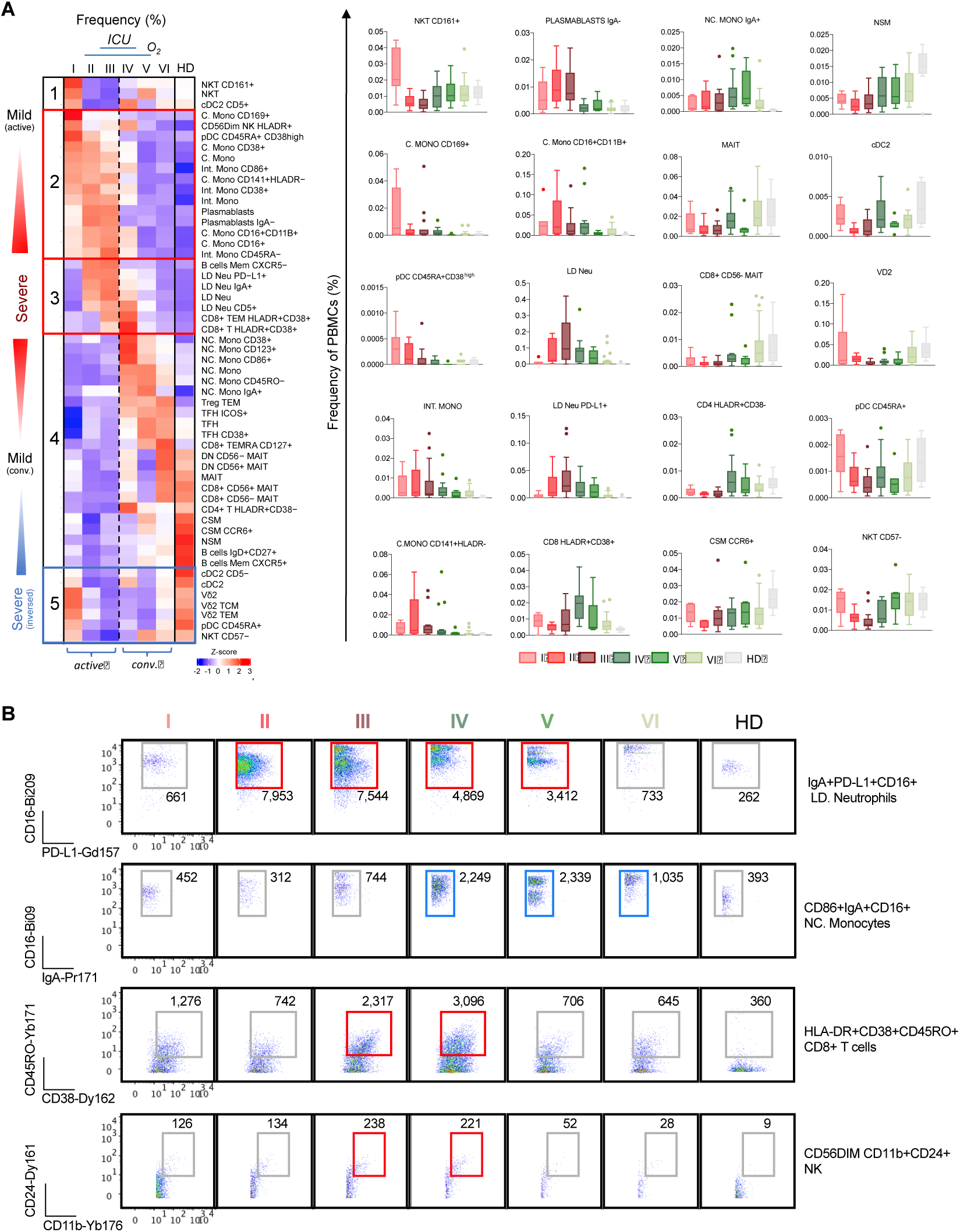
Association of immune cell types and immunotypes with disease severity. (A) Left: Heat map representation of frequencies of 53 immune cell populations with mild active, severe, and mild convalescence among the 6 group severity stratifications and divided into five clusters. Right: Box-and-whiskers plots showing means and IQR increased and reduced frequency of immune cell pollutions with disease severity. (B) Scatterplots of subsets of IgA^+^ LD Neu, CD86^+^ NC. Mono, HLA-DR^+^ CD8^+^ T and C56^Dim^ NK cells with each cell population additionally showing co-expression of CD16, PD-L1, IgA, CD45RO, CD38, CD24 and CD11b frequencies. The absolute number shown for the gated immune cells has been normalized per 300,000 PBMCs and thus reflects its frequency.

Among the T cells, the strongest association with severity was observed for HLADR^+^CD38^+^ CD8^+^ T cells. Unlike the frequencies of MAIT and NK cell subsets across groups I/II/III and groups IV/V/VI, which were largely invariant across groups and thus did not associate with severity (Figures S6A and S6B). The numbers of hyperactivated HLADR^+^CD38^+^ CD8^+^ T cells peaked in severity groups III/IV, whereby the highest frequency was detected in group IV during convalescence (Figures 4B and S7A). Also, a number of emerging studies suggested a role of inflammatory monocytes in the pathogenesis of COVID-19. In line with these results, although our data also showed elevated numbers of C. Mono and Int. Mono during the active phase, the increase with regard to severity was only marginal. The reverse trend was evident for NC. Mono, which declined in the active groups I/II/III but increased during convalescence (Figures 4B, 5A, S2A and S3D).

The association of the frequencies of the 38 basic immune subsets with the six severity groups as well as the associated surface marker expression is summarized in Figure S2. Their phenotypes can be assessed by using the online viewer (see Materials and Methods: Data and Code Availability). Figure 5A shows the subsets out of the 327 common immunotypes, which exhibited the most significant frequency changes with regard to the six severity groups. Based on the frequency distribution, they were divided into five different clusters (Figure 5A, left). Immune subsets in the first cluster positively associated with mild infection such as CD161^+^ NKT and CD5^+^ cDC2, which only increased in group I and may thus play a potential protective role (Figure 5A). The second cluster is positively associated with active SARS-CoV-2 infection across mild to severe groups I/II/II. Besides the plasmablasts, CD169^+^/CD38^+^ C. Mono and CD86^+^/CD45RA^-^/CD38^+^ Int. Mono but not total monocytes remained elevated in groups II/III. Also, like CD169^+^ C. Mono, both CD38^high^CD45RA^+^ pDC and HLA-DR^+^CD56^DIM^ NK peaked in group I and remained high in groups II/III above healthy levels. Notably, HLA-DR^Low^CD141^+^ C. Mono, which has been reported to be over-represented in ICU patients did not show a pronounced association with the most severe group III but was rather increased throughout the active phase (Silvin et al., 2020). This suggests that the aforementioned subset is unlikely to be a marker for severity. The third cluster is positively associated with severe COVID-19. In this cluster, CXCR5^+^ B cells memory subset was only increased in more severe groups II/III but various LD Neu subsets were elevated in both severity groups III and IV (Figures 5A and 5B). PD-L1^+^ LD Neu are most abundant in groups II/III while CD5^+^ LD Neu peaked in group IV. Within group IV, hyperactivated HLA-DR^+^CD38^+^ CD8^+^ T cell subset was also elevated (Figures 5A and,5B). The fourth cluster consisted of mixed frequency distribution during convalescence. Various NC. Mono subsets (CD38^+^/CD123^+^/CD86^+^/CD45RO^-^/IgA^+^) and Treg TEM and T_FH_ subsets were increased in frequencies across groups IV/V/VI above healthy levels (Figures 5A and S3G). On the other hand, immune subsets of MAIT, CSM and NSM of more severe COVID-19 patients often did not recover to healthy levels. Finally, the fifth cluster defined as “severe inverse” included subsets exhibiting the greatest loss in frequency in patients with most severe but not mild COVID-19. These were mostly cDC2, CD45RA^+^ pDC, CD57^-^ NKT and Vδ2 memory T cells, which may indicate direct involvement in COVID-19 related pathology (Figures 5A and S4).

The shifts in surface marker co-expression for some of the key populations are also shown in Figure 5B. For LD Neu, besides the IgA^+^ marker, the PD-L1 co-expression was particularly pronounced across the active and convalescent groups II to V. Additionally, CD16^high^ LD Neu was more abundantly expressed during convalescence, which was not seen in healthy donors (Figures 5B and S3C). Similarly for NC. Mono, besides IgA^+^ and CD86^+^ markers, CD16 co-expression was strongly increased in convalescent phase groups IV and V (Figures 5B, S3E and S3G). And for hyperactivated HLA-DR^+^CD38^+^ CD8^+^ T cells, the co-expression of CD45RO marker was highly elevated in groups III and IV. Similarly, a concomitant increase in the co-expression of CD11b^+^ and CD24^+^ markers by CD56^dim^NK was particular evident in groups III and IV (Figures 5B and S2B). Notably, this specific immunotype has been reported to transdifferentiate into myeloid cells e.g. neutrophils (Song et al., 2020).

### Dynamics of plasma cytokine levels in COVID-19 patients

In order to correlate the shifts in frequency and phenotype with the cytokine/chemokine environment, we have screened the corresponding COVID-19 plasma samples for 28 analytes using either Luminex or high-sensitivity Quanterix bead arrays. Of these, thirteen cytokines showed statistically significant associations with the six severity groups (adjusted p < 0.05) (Figure 6A). A comparison between the association with severity and the timing of the immune response revealed that IFNα, while abundantly found in the early active phase, had very little prognostic value with regard to severity (Figure 6B). While the levels of IL6 seem to better match disease severity, this applied only for the active phase. Also, IL6 was mostly detected during the early active phase. The association with severity however substantially improved for IP10, TNFα, HGF and VEGF-A. Especially the latter two were detected in similar extent during both the active and convalescent phases.

**Figure 6.**
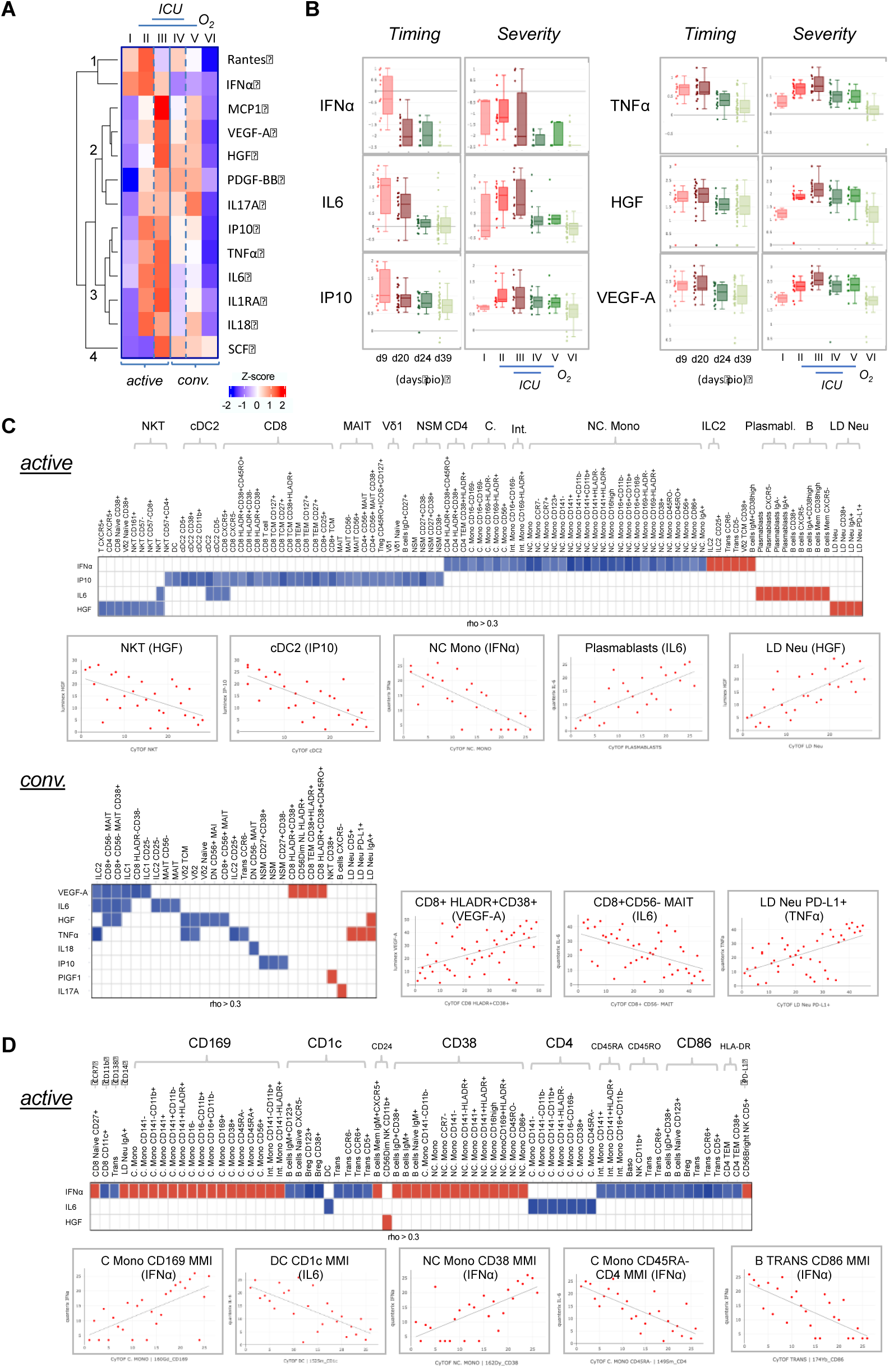
Characterization of cytokines in COVID-19 patients. (A) Changes in cytokine levels among COVID-19 patients based on timing and severity. The heatmap shows the z-scores of the mean logarithmically transformed concentration of the 13 cytokines (of a total of 28) showing significant differences between any of the 6 severity groups. The z-scores are colored in red for positive values and in blue for negative values. The cytokines are clustered using hierarchical clustering using Euclidean distances into four clusters, which are labeled 1 to 4 in the figure. (B) Box plots of selected cytokines showing differences in timing and/or severity. The timing (left panels) refers to the plasma cytokine levels detected on the respective day post illness onset (PIO), the severity (right panels) to the levels detected in the 6 severity groups. Red colors refer to samples from the active phase, green to convalescence phase. An interactive viewer is available in the online content: data availability section. (C) Associations between cytokine level and cell frequency during active and convalescent phase. The heatmap displays the strength of the association indicated by the correlation coefficient (rho). Color indicated the direction (Red: positive, blue: negative). Only associations abs(rho) > 0.3 and p < 0.05 are shown. Selected examples of these correlations are shown in the scatter plots. An interactive viewer is available in the online content: data availability section. (D) Associations between cytokine level and cell surface marker expression during the active phase. The plots are arranged as in Figure 6C except that the correlation was carried out with the MMI values of the various cell surface marker instead of the percentage value of the cell populations.

To further identify the cytokine/immune cell associations during SARS-CoV-2 infection, we first correlated the cytokine levels with the frequencies of the 327 immunotypes. The various plots for these correlations can be assessed in an interactive online viewer (see Materials and Methods: Data and Code Availability). As the immune responses in different phases of infection are vastly different, we carried out separate correlations for the active and convalescent phases (Figure 6C, top). During active infection, IFNα, IP10, IL6 and HGF showed a pleiotropic effect on a variety of immune cells. IFNα had a negative correlation with various monocyte subsets such as NC. Mono but positively associated with ILC2, CD38^+^ Vδ2 TCM, IgM^+^CD38^high^ B cells, CD5^-^ transitional B cells, CCR6^-^ transitional B cells and CD25^+^ ILCs. IL6 was positively associated with B cells and various plasmablast subsets, and HGF was positively associated with LD Neu but negatively with NKT.

During convalescence, a greater number of cytokines was associated with various immunotypes, albeit the association was typically weaker as compared to the active phase (Figure 6C, bottom). VEGF-A was strongly associated with HLA-DR^+^CD38^+^ CD8^+^ T cells but negatively with CD8^+^CD56^-^ MAIT cells. Interestingly, IgA^+^, CD5^+^ and PD-L1^+^ LD Neu positively associated with TNFα, which is known to promote neutrophil degranulation (Cross et al., 2008; Salamone et al., 2001). Additionally, PIGF1 and IL17A positively associated with CD38^+^ NKT and CXCR5^-^ B cells, respectively, while negative associations of both proinflammatory TNFα and HGF with Vδ2 T cells were observed, as well as negative associations of both IL6 and HGF with CD8^+^CD56^-^MAIT.

As cytokines can also directly influence the surface expression of biomarkers, we carried out a similar correlation for the surface molecules detected during the active phase on the 327 immunotypes (Figure 6D). While there were fewer cytokines found significant, they often correlated with the same surface marker on a number of different cell subsets. IFNα was positively associated with upregulated CD169^+^ on both C. and Int. Mono. The same cytokine also triggered the upregulation of CD38 on various NC. Mono as well as CD141^-^CD11b^-^ C. Mono and various B cell subsets. IFNα was also found negatively associated with CD86 on B cells, while IL6 was negatively associated with CD4 on various C. Mono populations. HGF was strongly associated only with the transdifferentiating CD11b^+^CD56^dim^ NK cells, which expressed CD24 (Song et al., 2020). Also, these data can be visualized using the online viewer (see Materials and Methods: Data and Code Availability).

### Integrated network analysis of immune subsets and plasma cytokines

Current data analysis so far has revealed a number of subsets and cytokines clearly linked to COVID-19 but the interaction of these components as well as the underlying pathways appear to be very complex and under-appreciated. To obtain a comprehensive overview of the interplay of these key players, we performed Bayesian network analysis using the complete sets of the mass cytometry data of the 327 cell subsets as well as the corresponding bead array measurements of the 13 cytokines (Figure 7 and Table S3). In order to separate the immune signatures of COVID-19 severity from those involved in anti-viral immune response, the fluctuations in these datasets were correlated independently with regard to the time point of sampling during the SARS-CoV-2 infection (Figures 7A and 7B) and the severity score assigned to the respective patient (Figures 7C and 7D). Interactive viewer for each of the networks is available as online resource (see Materials and Methods: Data and Code Availability)

**Figure 7.**
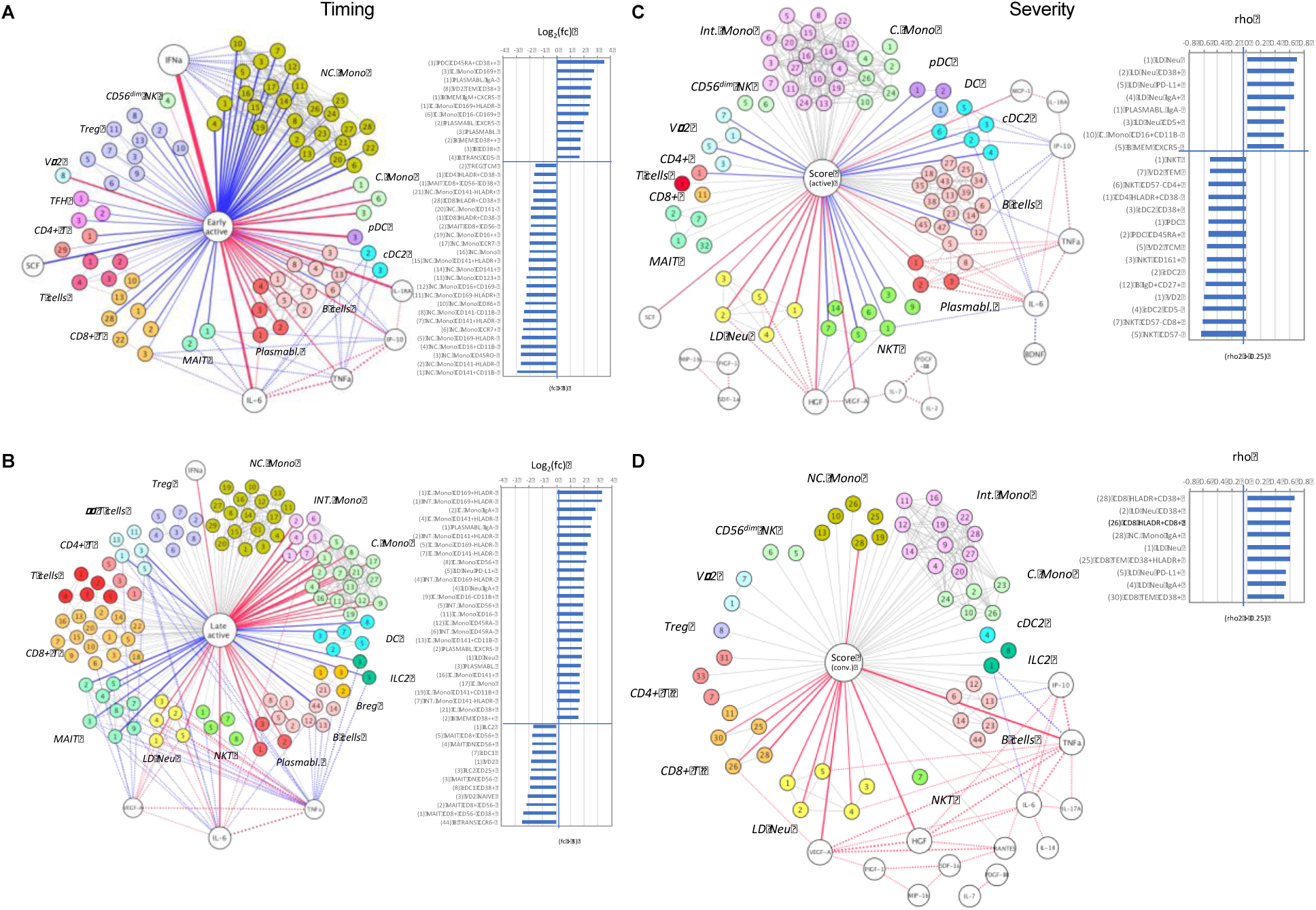
A node-edge interaction network of the cytokine level and immune cellular frequencies in COVID-19 patients. Association are shown with regard to the timing (A, B) and the severity (C, D). Nodes represent either cytokines (white) or immune subsets (colored). The central node represents the “comparison of interest”. The edges represent significant associations between two nodes with the thickness indicating the strength either based on fold change or correlation coefficient (rho). Color indicates the direction (Red: positive, blue: negative), dotted lines indicate associations with cytokines. For the central node, only associations with abs(rho) > 0.3 and p < 0.05 are colored and shown as bar charts on the right. For the timing (A, B) these bar charts indicate the fold changes in the early active (A) and late active state (B) in reference to late convalescent state while for the severity (C, D) they represent the correlation coefficient (rho) in reference to the severity groups in the active (C) and convalescent state (D). The number code of the cell subsets is listed in Table S3, an interactive network viewer is available in the Materials and Methods: data and code availability subsection and Key Resource Table. ^++^ denotes <u>high</u>ly stained immunotype.

For the timing of the immune response in the active phase of SARS-CoV-2 infection, the fold change of subset frequency and cytokine levels was determined independently for the median day 9 PIO (“early active”; Figure 7A) and median day 24 PIO (“late active”; Figure 7B). In the early active phase, the strongest correlation was observed for IFNα (Figure 7A). The increase in levels of IFNα inversely correlated with a general decline in the number of NC. Mono including the CD141^+^CD11b^-^, CD141^-^HLA-DR1^-^, CD45RO^-^ and CD16^+^CD11b^-^ subsets. For these cells, the absolute fold change was between 5 to 8, and the most pronounced reduction observed during the early active phase. The most substantial increase was detected for CD38^high^ CD45RA^+^ pDC and subsets of C. Mono expressing CD169, a marker directly induced by IFNα (compare Figure 6D). Other subsets positively associated with the early phase of infection included several subsets of plasmablasts, activated B memory and Vδ2 T cells. Of these, IgA^-^ plasmabasts positively correlated with IL6, which is also significantly upregulated in the early phase. The increase in plasma IL6 was also associated with a loss of CD8^+^ T cells, MAIT and cDC2. In addition, elevated IP10 was a node associated with a depletion of MAIT and certain B cell subsets (NSM CD27^+^CD38^-^ and NSM CD27^-^CD38^+^). However, we did not find any nodal association with Vδ2 T cells, C. Mono and pDC populations to any of the cytokines, while SCF and IL1RA were nonetheless early responders in the blood.

During the late active phase of SARS-CoV-2 infection, the IFNα levels dominating the early phase strongly subsided (Figure 7B). Although the correlation with NC. Mono nearly diminished, Int. and C. Mono strongly expanded in this phase, in particularly for the CD169^+^ subsets with a 8-fold increase in frequency. Next, the correlation with IL6 levels maintained at the late active phase but also two other cytokines, VEGF-A and TNFα, begun to show multiple cellular associations corresponding to significant losses of γδ T, ILC2 and MAIT cells, while IL6 was a node associated with late increase in LD Neu and Int. Mono, potentially linking these cytokines to the lymphopenia, neutrophilia and monocytosis, which are characteristic of SARS-CoV-2 infection.

A strikingly different picture emerged when the network analysis was carried out in reference to the severity of COVID-19 manifestation. Due to the different nature of the immune response during active infection and convalescence, two independent networks were generated for severity groups I to III (“active”; Figure 7C) and groups IV to VI (“conv.”; Figure 7D). The degree of association was determined by a linear correlation of the subset frequencies and cytokine levels in these groups and is expressed by their rho value. Notably during the active phase, the strongest positive correlation was observed for LD Neu and its CD38^+^, PD-L1^+^, IgA^+^ and CD5^+^ subsets (Figure 7C). With the exception of the latter, all of them also positively correlated with HGF, which was also directly associated with disease severity. While an increase in HGF correlated with an increased frequency of LD Neu subsets, it was also associated with a loss of NKT cells, namely of the CD57^-^ subsets. Although their frequencies were inversely associated with disease severity, this does not necessarily indicate a beneficial effect, as the disappearance of these subsets from the blood may likely be a consequence of their redistribution towards the inflamed tissue. Other subsets, which positively associated with severity are IgA^-^ plasmablasts and CD16^+^CD11b^-^ C. Mono, possibly pointing to the role of IgG in the COVID-19 pathogenesis. Moreover, plasmablast frequencies directly correlated with IL6 levels, which were also inversely associated with cDC2. Similar to pDC and Vδ2 T cells, their frequencies in the blood also inversely correlated with disease severity.

The direct association of the LD Neu subsets with severity was also observed during convalescence, and no significant negative correlation was detected for any of the immune subsets (Figure 7D). Positive correlations were found for IgA^+^ NC. Mono, several CD8^+^ TEM subsets, as well as CD38^+^, CD5^+^, IgA^+^ and PD-L1^+^ LD Neu. Of the cytokines, VEGF-A, HGF and TNFα correlated directly with severity, while fewer interactions between cytokines and cell subsets were detected. At least during convalescence, the increase in TNFα but not HGF was positively associated with IgA^+^ LD Neu and PD-L1^+^ LD Neu subsets (Cross et al., 2008). Additionally, elevated VEGF-A levels strongly associated with enriched HLA-DR^+^CD38^+^CD8^+^ T cells(Voron et al., 2015), which are also directly associated with severity. While especially the early response against SARS-CoV-2 seems to be dominated by IFNα- and IL6-driven immune responses (Figures 7A and 7B), network analysis did not provide any evidence that the pathways and cellular components activated by these cytokines have a major impact on the severity of the disease in either the active phase of infection (Figure 7C) or during convalescence (Figure 7D).

## Discussion

COVID-19 is triggered by viral infection whose pathology is mainly caused by collateral or even autoimmune-like damage inflicted by a hyperactivated immune system. By analyzing the blood samples of 77 hospitalized COVID-19 patients by mass cytometry and cytokine bead arrays, we attempted to delineate the key factors driving the immune pathology of COVID-19 from the genuine defense against SARS-CoV-2. To achieve this, we segregated the PBMCs into 327 distinct immune cell subsets and determined the plasma levels of 28 different chemokines and cytokines. The data were stratified with regard to different phases of the viral infection and correlated with the severity score defined by the clinical state of the patients. This generates a comprehensive data resource to analyze the immune response triggered by the SARS-CoV-2 infection in an integrated and interactive way.

A simplified timeline could be established by the breakdown of the infection into four phases (early/late active and early/late convalescence). The time point of the given cell subset was assessed by comparing their relative frequencies in these phases, and their contribution to severity was quantified by the rho^2^ parameter of the disease score correlation (Figure S8). Based on this, we propose the following immune progression: infection with SARS-CoV-2 leads to early IFN-α production, as evidenced by an early upregulation of CD169 on monocytes. This drives the disappearance of NC. Mono from the blood, which together with Vδ2 T, cDC2, pDC, NKT, CD8^+^ MAIT and CD8^+^ T cells appear to be recruited to the inflamed tissues. In parallel, during this early active phase (median days PIO: 9) there is also an expansion of activated memory B cells and importantly, antibody-producing plasmablasts. The plasmablast frequency remains high during the late active phase, a process associated with elevated levels of the proinflammatory cytokines IL6, IP10 and TNFα.

The late active phase (median days PIO: 20) is also characterized by an expansion of C. Mono, which likely differentiates into Int. and NC. Mono. The relative fraction of Int. Mono remained high during convalescence, above healthy controls even at the late convalescent phase (median days PIO: 39). In comparison, NC. Mono frequency was the highest during convalescence (Figure 3A). The same applies for CD8^+^ TEM cells, in particular for the HLADR^+^CD38^+^ subset, which has been found to be enriched in lungs of deceased COVID-19 patients (Xu et al., 2020). Notably, major changes in the frequency of NK subsets were only observed during convalescence, suggestive of a minor role during the early phase of infection.

Disease resolution and convalescence usually result in the normalization of most cell subsets. Notably, the late convalescent subsets, including hyperactivated HLADR^+^CD38^+^ CD8^+^ T cells, senescent CD57^+^ CD4^+^ T cells, Treg TCM/TEM, activated ICOS^+^/CD38^+^ T_FH_, NC. Mono, Vδ2 TCM/TEM, lymph node homing CCR6^+^ CSM B cells, NSM B cells and IL12-producing CD5^-^ cDC2 cells, persistently failed to recover to healthy levels, and may contribute to the lingering symptoms associated to post-infection aberrations (Figure 3B). Further studies would be needed to determine if alterations to these immune subsets contribute to the long-term effects observed in some patients after recovery from severe disease (Weerahandi et al., 2020).

While the involvement of immune cell subsets in anti-viral defense can be heterogeneous, a clearer picture emerges when considering specific immunotypes related to COVID-19 severity. Among the frequencies of subsets such as NC. Mono, Vδ2 T, cDC2, pDC, CD57^-^ NKT, CD8^+^ MAIT, and CD8^+^ T cells showing depletion or enrichment for plasmablasts during infection (Figures S3-S7), the most dramatic severity association was observed for LD Neu (Figure S3). Although LD Neu is virtually absent in the blood of healthy donors, extreme levels were often detected in the blood in the most severe group of patients requiring supplemental O_2_ and/or ICU. Due to their high density, neutrophils are not expected to be part of the PBMC fraction. However, processes like neutrophil degranulation can reduce the cell density, which allows them to ‘contaminate’ the lymphocyte/monocyte fraction during Ficoll separation (Hacbarth and Kajdacsy-Balla, 1986). As degranulation is an integral part of neutrophil biology, the number of LD Neu in the blood may indicate the extent of ongoing neutrophil responses. Previous studies have reported increased numbers of activated neutrophils inside the inflamed lung tissue (Barnes et al., 2020; Fox et al., 2020; Wang et al., 2020; Yao et al., 2020) as well as elevated levels of immature neutrophils in the blood circulation of COVID-19 patients (Carissimo et al., 2020). This is in line with another recent study showing the presence of neutrophil extracellular traps (NETs) in the lungs of deceased COVID-19 patients (Radermecker et al., 2020). Notably, LD Neu seems to have a heightened capacity to release NETs (Yu and Su, 2013). In fact, LD Neu and NET formation have been reported in a number of autoimmune diseases, such as antiphospholipid syndrome (Mauracher et al., 2020), systemic lupus erythematosus(Van Den Hoogen et al., 2020), and anti-neutrophil cytoplasm autoantibody vasculitis (Ui Mhaonaigh et al., 2019), potentially consistent with autoimmune-like features of COVID-19 pathology (Woodruff et al., 2020).

As monocytes has been implicated in COVID-19 pathology (Bedin et al., 2020; Guo et al., 2020; Silvin et al., 2020; Zhou et al., 2020), we further characterized the monocyte population into about 90 different subsets. As expected, IFNα is linked to the induction of CD169^+^ C. Mono during the early active phase as well as the expansion of CD16^hi^ monocyte populations, however we failed to observe any significant association with disease severity. The only exception may be the expansion of CD16^+^CD11b^-^ C. Mono (Figure 7C) during the active phase and possibly IgA^+^ NC. Mono during convalescence (Figure 7D). Here, we could neither confirm the protective effect of CD169^+^ C. Mono nor the increase of CD141^+^HLADR^-^ C Mono in severe cases as reported by Hadjadj et al. (Hadjadj et al., 2020). A possible explanation for the discrepancy may be the difference in the timeline of sample taking or in the respective definition of severity; in Hadjadi’s study, ‘mild’ cases were essentially asymptomatic but all of our patients were hospitalized.

Lastly, our study goes a step further by directly linking the neutrophil-specific immune response to the pathology of COVID-19. One of the most striking results is the strong association between LD Neu frequency and HGF plasma levels. Both factors are directly associated with COVID-19 severity, suggesting that the cytokine may also play a key role in the pathology. The associative nature of this study however does not allow us to draw direct causal conclusions. Furthermore, the source of HGF is still controversial. While the bulk of HGF seems to be released by mesenchymal cells, neutrophils may still play an important role by acting as a source of matured HGF (Ohnishi et al., 2006; Small and Lung, 2006; Wislez et al., 2003). HGF typically acts as anti-inflammatory and supports wound healing. For instance, mesenchymal stem cells have been shown to alleviate acute lung injury via the paracrine secretion of HGF (Lu et al., 2019). HGF and its receptor c-MET also play a crucial role in various cancer where the neutrophil/HGF axis seems to mediate tumor growth by eliciting immune-suppression (Glodde et al., 2017). However, depending on differentiation and tissue environment, neutrophils can exert both pro- and anti-inflammatory effects (Rosales, 2018). An earlier *in vitro* study even suggested that HGF stimulates neutrophil degranulation (Kowanko et al., 1993). It is therefore likely that the presence of HGF in severe COVID-19 may also promote the release of NETs (Radermecker et al., 2020). It may thus be worth to evaluate existing HGF/c-MET drugs with regard to their ability to prevent or stop deteriorating COVID-19 symptoms.

In summary, while the cellular and molecular interactions underlying COVID-19 are very complex, we have identified some new candidates for potential treatment interventions. Here, the HGF/LD Neu axis may represent a promising new lead for direct interventions, and future studies have to show if HGF is actually a better pharmacological target than IFNα and IL6. The trials with IL6 inhibitors have already failed and there is acute demand for effective treatments. In support of this hunt, this study generates a database that could be used as a crucial resource to evaluate the pathways and to validate, identify or exclude candidate targets that could help to control the ongoing pandemic.

## Materials and Methods

### Study design, sample size and participants

For this study, 77 COVID-19 patients and 10 healthy donors were recruited. Enrollment of COVID-19 patients was via PROTECT, a Singapore COVID-19 cohort study among seven public health institutions. Healthy individuals were recruited under a Singapore Immunology Network study entitled, “Study of blood cell subsets and their products in models of infection, inflammation and immune regulation”. Both studies had received prior approval from their respective institutional review boards (IRBs). All individuals involved in this study were over the age of 21, comprising 66 males and 21 females. Additional demographic details can be found in Table S1.

### Sample collection

Blood from healthy adult donors and COVID-19 patients were collected in BD Vacutainer CPT Tubes and processed according to manufacturer’s instructions to obtain the PBMC and plasma fractions. Isolated PBMCs were then used for mass cytometry staining after two washes with 1X phosphate buffer saline (PBS).

### Cytometry by time-of-flight (CyTOF) sample processing and data acquisition

Freshly isolated ficoll-density centrifuged PBMCs were plated at 0.5 – 1 x 10^6^ in a 96-well V bottom plates and stained for viability with 100 µL of 66 µM of cisplatin (Sigma-Aldrich) for 5 minutes on ice. Cells were then washed with staining buffer (4% v/v fetal bovine serum, 0.05% v/v sodium azide in 1X PBS) and stained with anti-γδTCR-PE and anti-Vδ1-FITC in 50 µL reaction volume for 15 minutes at room temperature. Cells were washed with staining buffer and then stained with 50 µL of metal isotope-labeled surface antibodies on ice. After 20 minutes, cells were washed with staining buffer, followed by PBS, and fixed in 4% v/v paraformaldehyde (PFA, Electron Microscopy Sciences) at 4°C overnight. On the following day, cells were incubated in staining buffer for 5 minutes. Cellular DNA was labeled at room temperature with 170 nM iridium intercalator (Fluidigm) in 2% v/v PFA/PBS. After 20 minutes, cells were washed twice with staining buffer.

Prior to CyTOF acquisition, cells were washed twice with water before final re-suspension in water. Cells were enumerated, filtered and diluted to a final concentration of 0.6 x 10^6^ cells/mL. EQ Four Element Calibration Beads (Fluidigm) were added to the samples at a final concentration of 2% v/v prior to acquisition. Samples were acquired on a Helios Mass Cytometer (Fluidigm) at an event rate of < 500 events per second. After CyTOF acquisition, data were exported in flow-cytometry (FCS) format, normalized to 300,000 PBMCs and events with parameters having zero values were randomized using a uniform distribution of values between minus-one and zero. Subsequently, manual gating was performed to exclude residual beads, debris and dead cells.

### Gating strategy for CyTOF

We have designed a 40-plex antibodies panel for mass cytometry and performed non-supervised Uniform Manifold Approximation and Projection (UMAP) or Triplet-constraint (TriMAP) dimensionality reduction for larger dataset embedding of ficoll-density centrifuged PBMCs obtained from both COVID-19 active and convalescent patients (Amid and Warmuth, 2019; McInnes et al., 2018). Iterative manual and UMAP clustering identified populations of T cells, B cells, monocytes, NK, DC, ILCs, basophil as well as the LD neutrophils based on their cell surface expression markers to generate 327 different immune cell subpopulations.

### Multiplex microbead-based Luminex immunoassays

Plasma samples were treated by solvent/detergent based on Triton™ X-100 (1%) for virus inactivation (Darnell and Taylor, 2006). Immune mediator levels in COVID-19 patient plasma across different active and convalescent groups were measured with 24-plex Human ProcartaPlex™ (ThermoFisher Scientific). The kit analyte detection panel included brain-derived neurotrophic factor (BDNF), beta-nerve growth factor (bNGF), hepatocyte growth factor (HGF), monocyte chemoattractant protein (MCP) 1, macrophage inflammatory protein (MIP) 1α, MIP1β, RANTES (regulated on activation, normal T cell expressed and secreted), stromal cell-derived factor 1 (SDF1α), interferon (IFN) gamma-induced protein 10 (IP10), IFNγ, interleukin (IL) IL1β, IL1RA, IL2, IL5, IL6, IL7, IL18, IL12p70, leukemia inhibitory factor (LIF), stem cell factor (SCF), tumor necrosis factor (TNFα), vascular endothelial growth factor A (VEGF-A), platelet derived growth factor (PDGF-BB), and placental growth factor (PLGF1).

Plasma from COVID-19 patients, healthy controls, as well as standards were incubated with fluorescent-coded magnetic beads pre-coated with respective antibodies in a black 96-well clear-bottom plate overnight at 4°C. After incubation, plates were washed 5 times with wash buffer (PBS with 1% v/v bovine serum albumin (Capricorn Scientific) and 0.05% v/v Tween-20 (Promega)). Sample-antibody-bead complexes were incubated with biotinylated detection antibodies for 1 hour and washed 5 times with wash buffer. Subsequently, Streptavidin-PE was added and incubated for another 30 minutes. Plates were washed 5 times again, before sample-antibody-bead complexes were re-suspended in sheath fluid for acquisition on the FLEXMAP® 3D (Luminex) using xPONENT® 4.0 (Luminex) software. Data analysis was done on Bio-Plex Manager™ 6.1.1 (Bio-Rad). Standard curves were generated with a 5-PL (5-parameter logistic) algorithm, reporting values for both mean florescence intensity (MFI) and concentration data.

Internal control samples were included in each plate to remove any potential plate effects. Readouts of these samples were then used to normalize the assayed plates. A correction factor was obtained from the median concentration values observed across the multiple assay plates and this correction factor was then used to normalize all the samples. The concentrations were logarithmically transformed to ensure normality. Analytes that were not detectable in patient samples were assigned the value of logarithmic transformation of the Limit of Quantification (LOQ).

### Multiplex microbead-based Quanterix immunoassays

Plasma immune mediator levels in selected active and convalescence phase of COVID-19 patients were measured using SIMOA Cytokine 3-Plex B (C3PB) assay kit (Quanterix) and SIMOA IFN-a assay kit (Quanterix). C3PB kit analyte detection included interleukin (IL) IL6, IL17A and tumor necrosis factor α (TNFα).

Standards and plasma from COVID-19 patients and healthy controls were pre-diluted in a 96-well plate before loading into the Simoa® HD-1 Analyzer (Quanterix) for data acquisition. Reagents from the C3PB and IFNα assay kits were prepared according to the kit manual and loaded into the analyzer. Fully automated data acquisition was done on Simoa® HD-1 Analyzer (Quanterix). Standard curves were generated with a 4-PL (4-parameter logistic) algorithm, reporting values for concentration data.

### Quantification and Statistical Analysis

Active and convalescence phase samples were defined by PCR positivity and serve as time based clinical end points. Active phase samples were further divided into early (PIO <= 14 days) and late (PIO > 14 days). Convalescence phase samples were also further divided into early (PIO <= 28 days) and late (PIO > 28 days). These provide a more granular time based clinical end points.

Severity based clinical end points were defined for active and convalescence phase samples separately. Three severity groups were defined for each phase consisting of symptomatic patients, patients requiring oxygen supplementation and patients requiring oxygen supplementation and awarded into intensive care unit as shown in Figure 1A.

Mass cytometry and cytokine measurements were associated to the clinical end points (time based as well as severity based) using Kruskal-Wallis tests followed by Dunn’s post hoc tests. Correlations between mass cytometry and cytokine measurements were done using Spearman Rank correlations. In the event that multiple samples from the same patient was available for same time period, the earliest of the samples were used for analyses to ensure that all samples used in the analyses are distinct. Multiple testing correction was done using the method of Benjamini and Hochberg. P values less than 0.05 were deemed to be significant. All statistical tests were two-sided (when appropriate) unless otherwise indicated. Statistical analyses were done using the R statistical language version 3.6.2. All statistical details are provided in the interactive viewers provided at https://data.mendeley.com/datasets/467s57xj8s/draft?a=15341765-e712-4eec-8107-a1d9c8da331a

Overviews of the mass cytometry immune cell subpopulations were generated using UMAP in R version 3.6.2 using the uwot package. Heat maps were generated in R version 3.6.2 using the CompexHeatmap package. Graphs of the significant associations were generated in R version 3.6.2 using the iGraph package and visualized in Cytoscape version 3.8.0. Additional visualizations were done in TIBCO Spotfire.

## Data and Code Availability

Data generated and/or analyzed during this study are available in the following public repositories and also at https://data.mendeley.com/datasets/467s57xj8s/draft?a=15341765-e712-4eec-8107-a1d9c8da331a.

An interactive viewer of the mass cytometry data associations with clinical endpoints (Figures 2A, 3, 5A and 5B and S2) are available at https://www.dropbox.com/s/wz93vwn2vvjsjry/cytof_sample_group_association_results_paper_vis_covid19_cytof_results_viewer.html?dl=1

An interactive viewer of the cytokine data associations with clinical endpoints (Figures 6A and 6B) are available at https://www.dropbox.com/s/4v107l3b65h5qfh/luminex_sample_group_association_results_vis_covid19_cytof_results_viewer.html?dl=1

The mass cytometry, cytokine and clinical data is available as an Excel file at https://www.dropbox.com/s/yd2spn3lholhuv3/all_cytof_multimodal_data_paper.xlsx?dl=1

An interactive viewer of interaction network in Figure 7A is available at https://www.dropbox.com/s/zm7a4s6nqelfnso/network_data_early_active_late_con_all_subset_percent_only_vis_bivariatetests.html?dl=1

An interactive viewer of interaction network in Figure 7B is available at https://www.dropbox.com/s/yt8sf6uwhtte55y/network_data_late_active_late_con_all_subset_percent_only_vis_bivariatetests.html?dl=1

An interactive viewer of interaction network in Figure 7C is available at https://www.dropbox.com/s/50tfhif6eqz3uoa/network_data_active_all_subset_percent_only_icu_regression_vis_bivariatetests.html?dl=1

An interactive viewer of interaction network in Figure 7D is available at https://www.dropbox.com/s/q43bz8s294bwobu/network_data_con_all_subset_percent_only_icu_regression_vis_bivariatetests.html?dl=1

An interactive viewer of cytokine data correlation with mass cytometry data at active phase (Figures 6C and 6D) is available at https://www.dropbox.com/s/6y6bo7zl40qlm26/luminex_correlation_analysis_active_convalescence_group_active_results_vis_stats_results_viewer.html?dl=1

An interactive viewer of cytokine data correlation with mass cytometry data at convalescence phase (Figure 6C) is available at https://www.dropbox.com/s/2h0awk6l4rgzmn9/luminex_correlation_analysis_active_convalescence_group_convalescence_results_vis_stats_results_viewer.html?dl=1

## Acknowledgements

This study was funded by grants from Singapore’s National Medical Research Council (NMRC)’s COVID-19 Research Fund (grant numbers COVID19RF-001, COVID19RF-004, and COVID19RF-007), as well as the Biomedical Research Council COVID-19 Fund (grant number H20/04/g1/006) and the A*ccelerate GAP Fund (grant number ACCL/19-GAP064-R20H-H) from the Agency of Science, Technology, and Research, Singapore. We thank Etienne Raimondeau from LaPipette for the design of Figure S8. We thank Mark I-Cheng Chen from the National University of Singapore and National University Health System for his support and patient cohort. We thank all clinical and nursing staff who provided medical care to the patients, staff at the Communicable Diseases Division of the Ministry of Health, Singapore, who contributed to the outbreak response and contact tracing, and staff at the Singapore Infectious Disease Clinical Research Network and Infectious Disease Research and Training Office of the National Centre for Infectious Diseases for coordinating patient recruitment.

## Author Contributions

Conceptualization: J.L., P.K.J., L.F.P.N., B.L., O.R.; Methodology: J.L., K.W.W.T., C.Y.L., K.P.T., G.C., Y.H.C., C.M.P., C.Y-P.L., S.-W.F., N.K.-W.Y., R.S.-L.C., S.N.A., Z.W.C., M.Z.T., A.T.-R., N.L.F., W.H.; Software: K.D., B.L.; Validation: J.L., P.K.J.; Formal Analysis: J.L., P.K.J., L.W.W., K.D., B.L.; O.R.; Resources: M.I.-C.C., S.-Y.T., L.Y.A.C., S.K., T.S.-Y., D.C.L., Y.-S.L., S.W.X.O., B.E.Y.; Data Curation: K.D., B.L.; Writing – Original Draft: J.L., P.K.J, L.W.W., B.L., O.R.; Writing – Review & Editing: J.L., P.K.J., L.W.W., B.L., O.R.; Visualization: J.L., P.K.J., L.W.W., B.L., O.R.; Supervision: L.R., L.F.P.N., O.R.; Project Administration: L.F.P.N., O.R.; Funding Acquisition: L.F.P.N.

## Competing Interests

The authors declare no competing interests

## Supplemental Information

**Figure S1.**
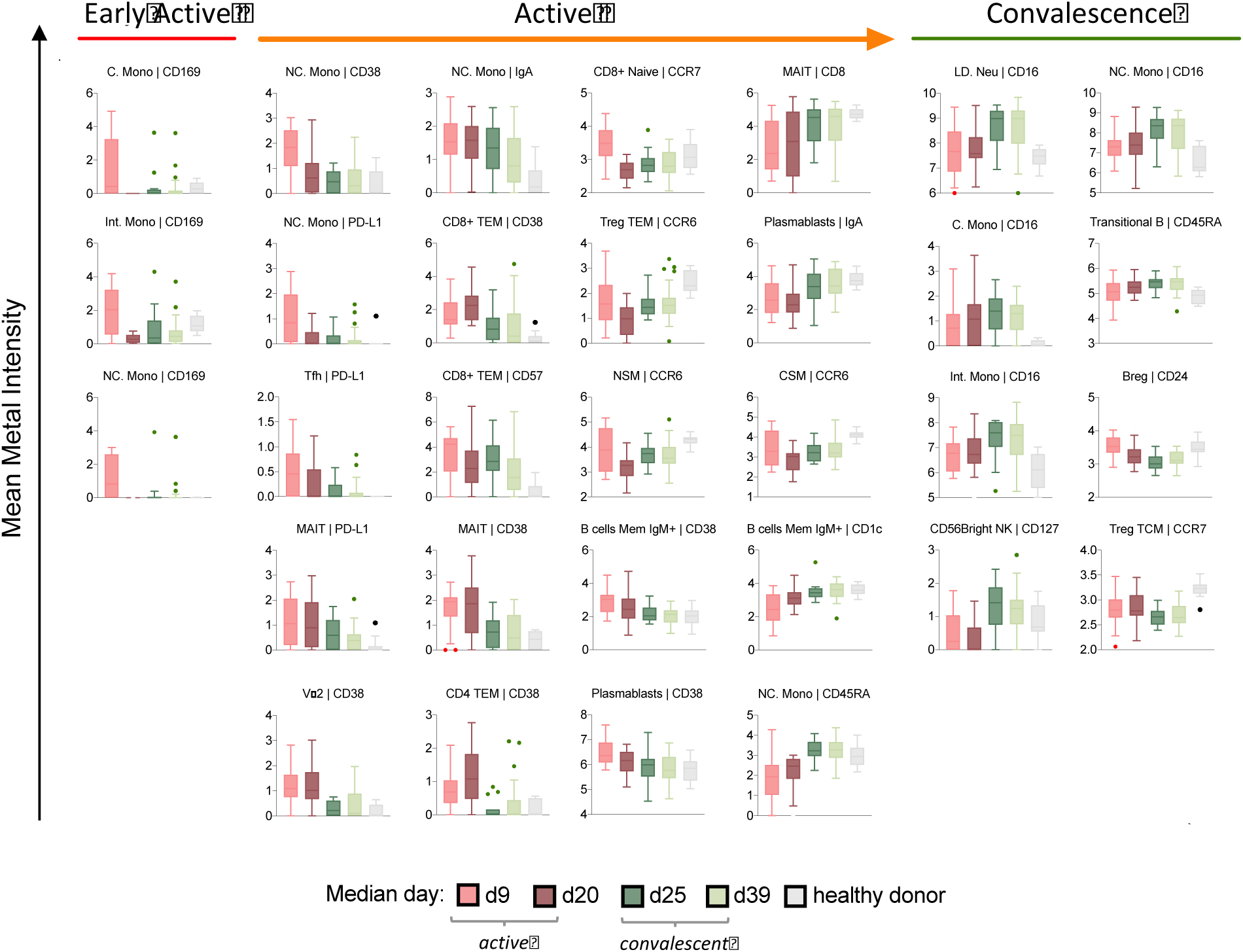
Temporal changes in surface marker expression profiles of various immune cell subsets during active and convalescent COVID-19.

**Figure S2.**
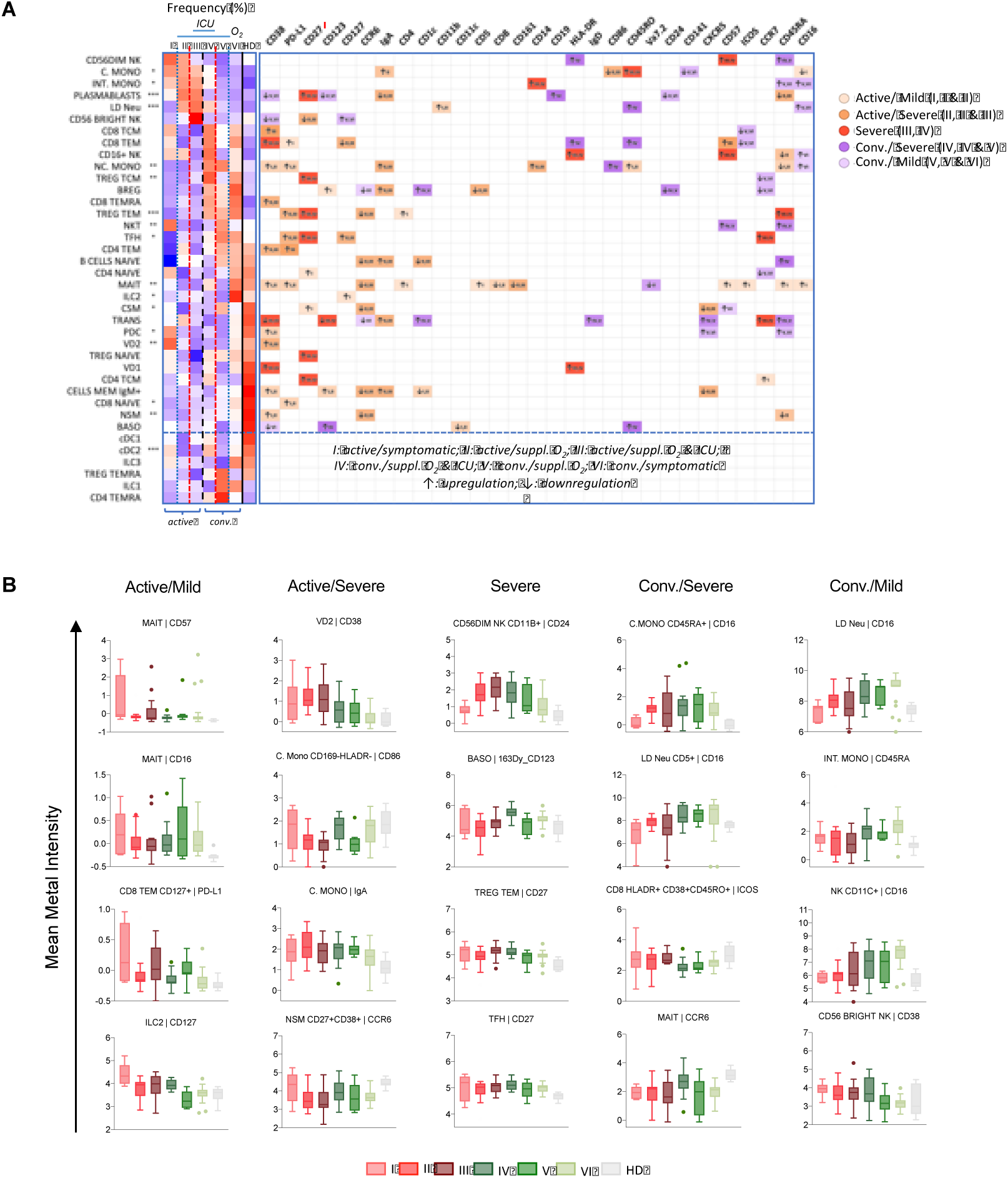
Alterations of immune cell types dynamics and immunotypes associated with mild and severe during active and convalescent infection. (A) Left: Heat map representation of frequencies of 38 main immune cell populations among the 6 groups severity stratifications. Right: up- or down-regulation of indicated surface markers for the 38 main immune cell populations among 6 group severity stratifications. Asterisks indicate statistical significance - *, p<0.05; **, p<0.01; ***, p<0.001 (one-way ANOVA of all disease phases and healthy controls). (B) Box-and-whiskers plots of selected immunotypes showing means and IQR up-regulation and down-regulation of surface markers associated with active/mild, active/severe, severe, conv./severe, and conv./mild clinical states.

**Figure S3.**
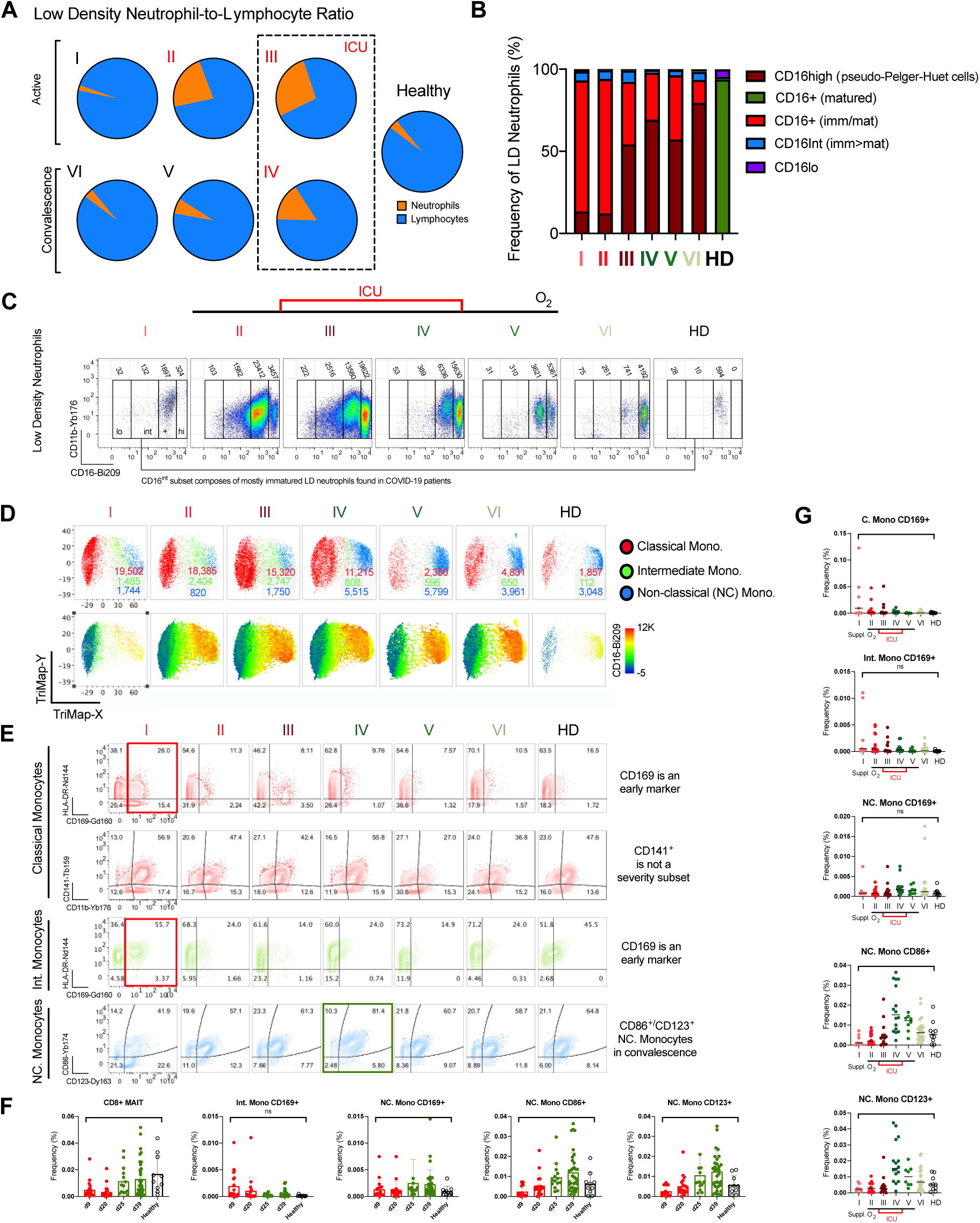
Neutrocytosis and monocytosis during SARS-CoV-2 infection. (A) Neutrocytosis due to SARS-CoV-2 infection is mild among symptomatic group I patients but persisted in severe convalescence group. A higher Low-Density Neutrophil-to-Lymphocyte Ratio is associated with disease severity. (B) Frequency distribution of Low-Density Neutrophil subsets whereby the CD16^+^ LD neutrophils are mostly mature neutrophils in healthy donors but co-mixed with immature neutrophils from the CD16^int^ LD neutrophil subset in COVID-19 patients. The presence of CD16^++/high^ LD neutrophils described as pseudo-Pelger-Huet cells, are absent in healthy donors. (C) The pseudo-coloured plots of LD neutrophil gated as CD16^lo^, left shift CD16^int^, CD16^+^ and CD16^high^ subsets and their indicated cell count out of 210K human PBMCs. (D) Monocytosis is apparent even in symptomatic group I and decreases with convalescence with CD16 marker upregulation in NC. Monocytes. (E) Comparison of NC. monocytes and other monocytes associated with convalescence and COVID-19 disease severity. Scatter plots depict the means with SEM. ns: not significant, *, p <0.03; **, p <0.002; ***, p <0.0002, ****, p <0.0001 (Kruskal-Wallis test with multiple comparisons performed on each disease severity group versus total healthy). (F-G) Frequency changes of representing monocyte subsets associated with surface markers during (F) SARS-CoV-2 infection and (G) COVID-19 severity.

**Figure S4.**
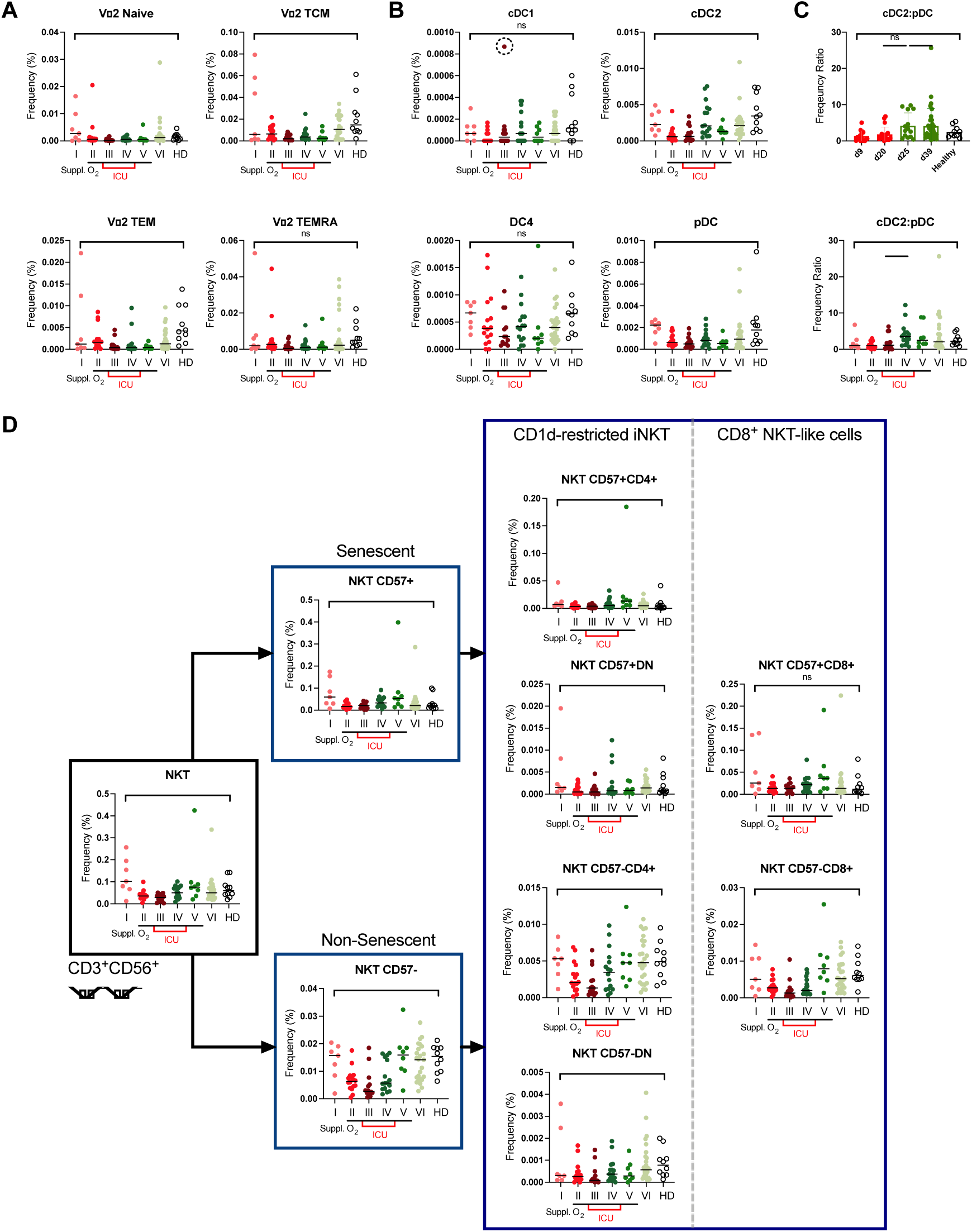
COVID-19 severity is specific to selected Vδ2 T, dendritic cell and non-senescent NKT cell subsets. Overview of frequency changes across the (A) Vδ2 T cells and (B) dendritic cells showing insignificance of subsets selected based on p-values. Scatter plots depict the means with SEM. ns: not significant,*, p <0.03; **, p <0.002; ***, p <0.0002, ****, p <0.0001 (Kruskal-Wallis test with multiple comparisons performed on each disease severity group versus total healthy). An extreme outlier due to a single patient in cDC1 is encircled. Vδ1 T cell subsets are insignificant and thus not shown. (C) The ratio of cDC2: pDC frequency during COVID-19 infection. (D) Clustering of NKT gated based on CD3^+^CD56^+^Vδ1^-^Vδ2^-^ T cell subsets into invariant NKT (iNKT) and CD1d-unrestricted CD8+ NKT-like cells gated based on senescent marker CD57. The three human NKT subsets (CD4^+^, CD8^+^ and CD4^-^CD8^-^/DN) also consist of Type I presenting invariant Vα24Jα18 TCR and Type II presenting a diverse TCR repertoire, which are undefined in this work.

**Figure S5.**
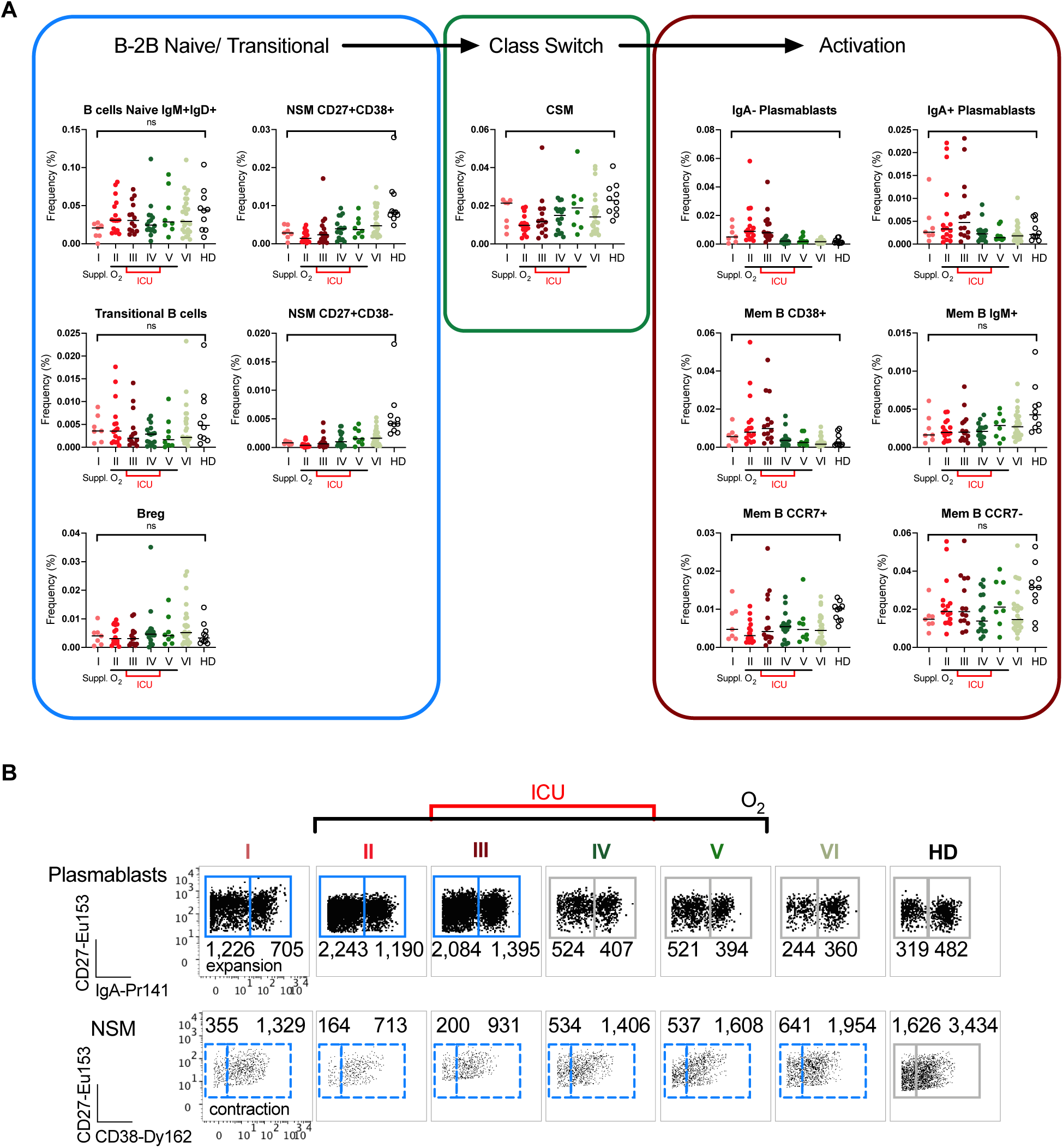
Heterogeneous B cell subset development during SARS-CoV-2 infection. (A) Scatterplots of B cell subset frequencies from Naïve to Memory based on COVID-19 severity. Scatterplots depict the means with SEM. ns: not significant, *, p <0.03; **, p <0.002; ***, p <0.0002, ****, p <0.0001 (Kruskal-Wallis test with multiple comparisons performed on each disease severity group versus total healthy). (B) Dotplots of representative B cell subsets for either expanded or contracted cell population in response to SARS-CoV-2 infection. The absolute numbers of cells out of 300,000 PBMCs are shown.

**Figure S6.**
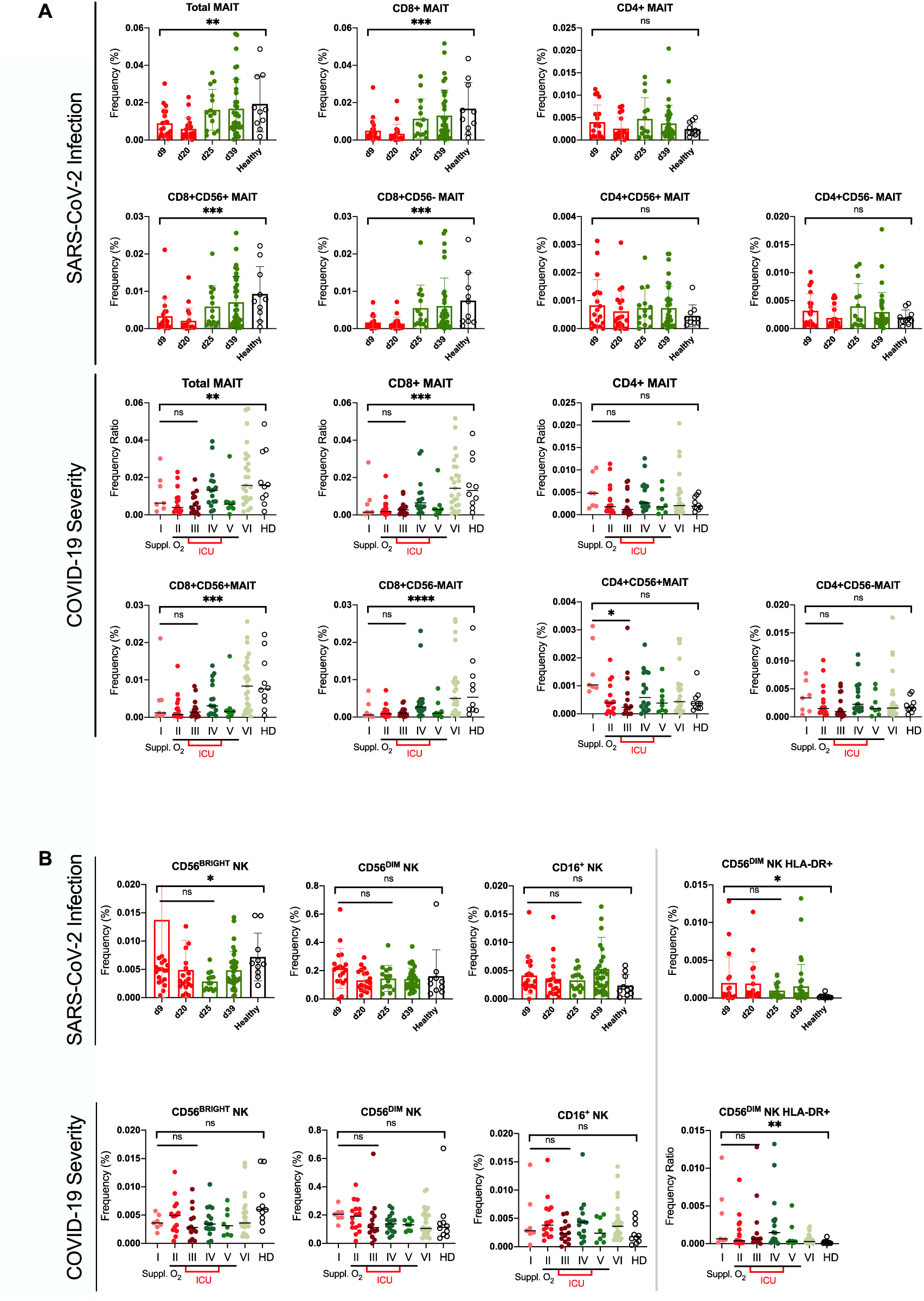
Loss of CD8^+^ MAIT and CD56^Bright^ NK cells are innate-like responses to SARS-CoV-2 infection but not severity. (A) Frequencies of MAIT cell subsets based on disease stage from early active to late convalescence. CD8^+^ but not CD4^+^ MAIT cells are significantly reduced during SARS-CoV-2 infection, which recover with health. Also, frequencies of different MAIT cell subsets with disease severity. There is little or no significance among severity groups I, II and III. (B) The depletion of CD56^Bright^ NK subpopulation when compared to healthy does not associate with COVID-19 disease severity. Also, CD56^Dim^ NK HLA-DR^+^ subset is not strongly correlated to disease severity. Scatter plots depict the means with SEM. ns: not significant, *, p <0.03; **, p <0.002; ***, p <0.0002, ****, p <0.0001 (Kruskal-Wallis test with multiple comparisons performed on each disease severity group versus total healthy).

**Figure S7.**
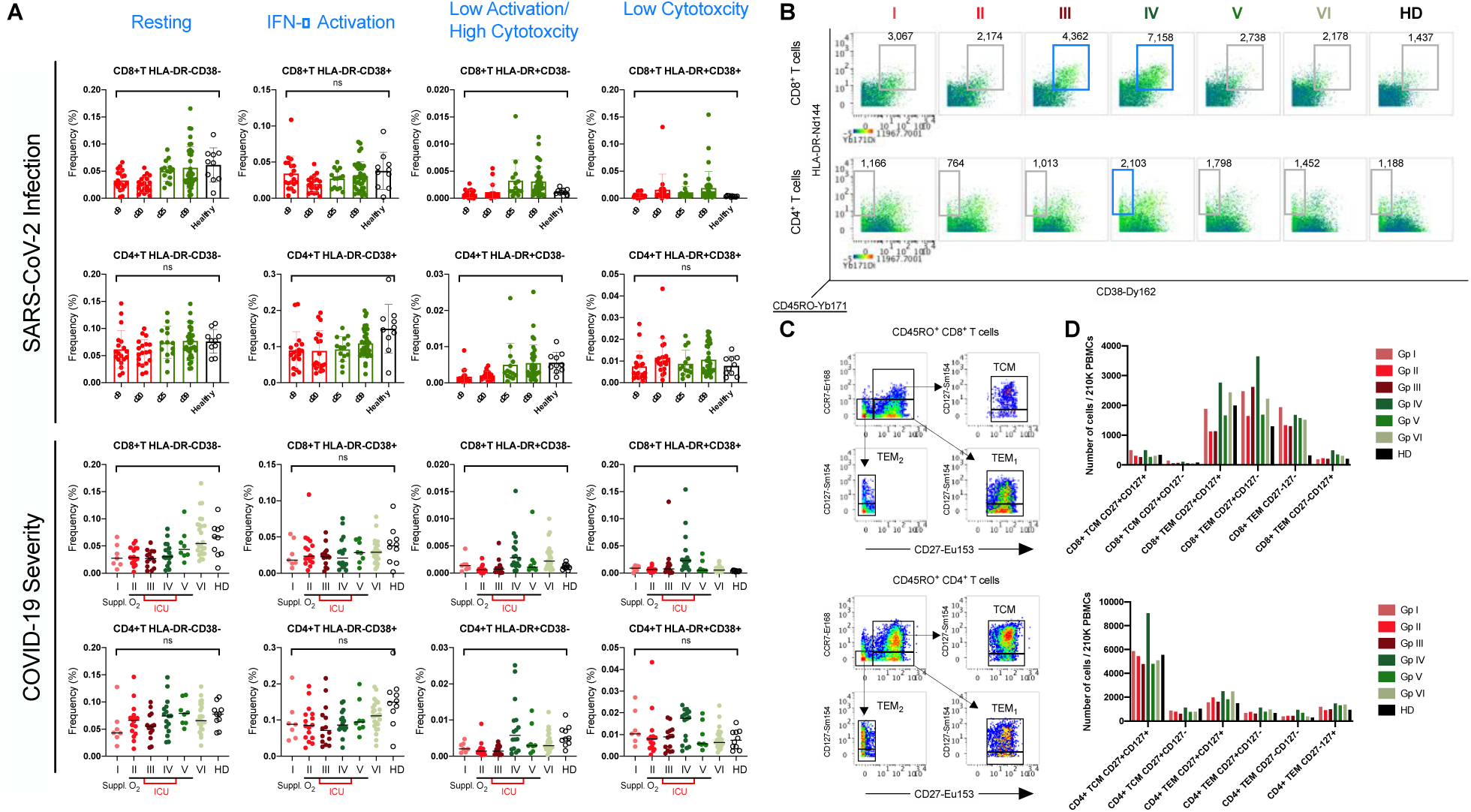
Heterogeneous T cell activation during SARS-CoV-2 infection. (A) Frequencies of CD4^+^ and CD8^+^ T cells based on HLA-DR and CD38 cell surface markers. Lymphopenia is apparent during active SARS-CoV-2 infection and T lymphocyte number increases in convalescence. Disease severity grouping further delineates finer structures. The CD8^+^ T cell subpopulations e.g. HLA-DR^+^CD38^+^ CD8^+^ T subsets are elevated in COVID-19 severity group IV. Scatter plots depict the means with SEM. ns: not significant, *, p <0.03; **, p <0.002; ***, p <0.0002, ****, p <0.0001 (Kruskal-Wallis test with multiple comparisons performed on each disease severity group versus total healthy). (B) 3-dimensional dotplots of statistically significant HLA-DR^+^CD38^+^ CD8^+^ and HLA-DR^+^CD38^-^ CD4^+^ T subset against memory CD45RO^+^ antigen. (C) Gating strategies for memory T subsets. (D) Barplot of total memory CD8^+^/CD4^+^ T subsets (TCM and TEM) and the CD27 and CD127 surface markers across disease severity groups showing similarities and differences among COVID-19 patients and healthy donors.

**Figure S8.**
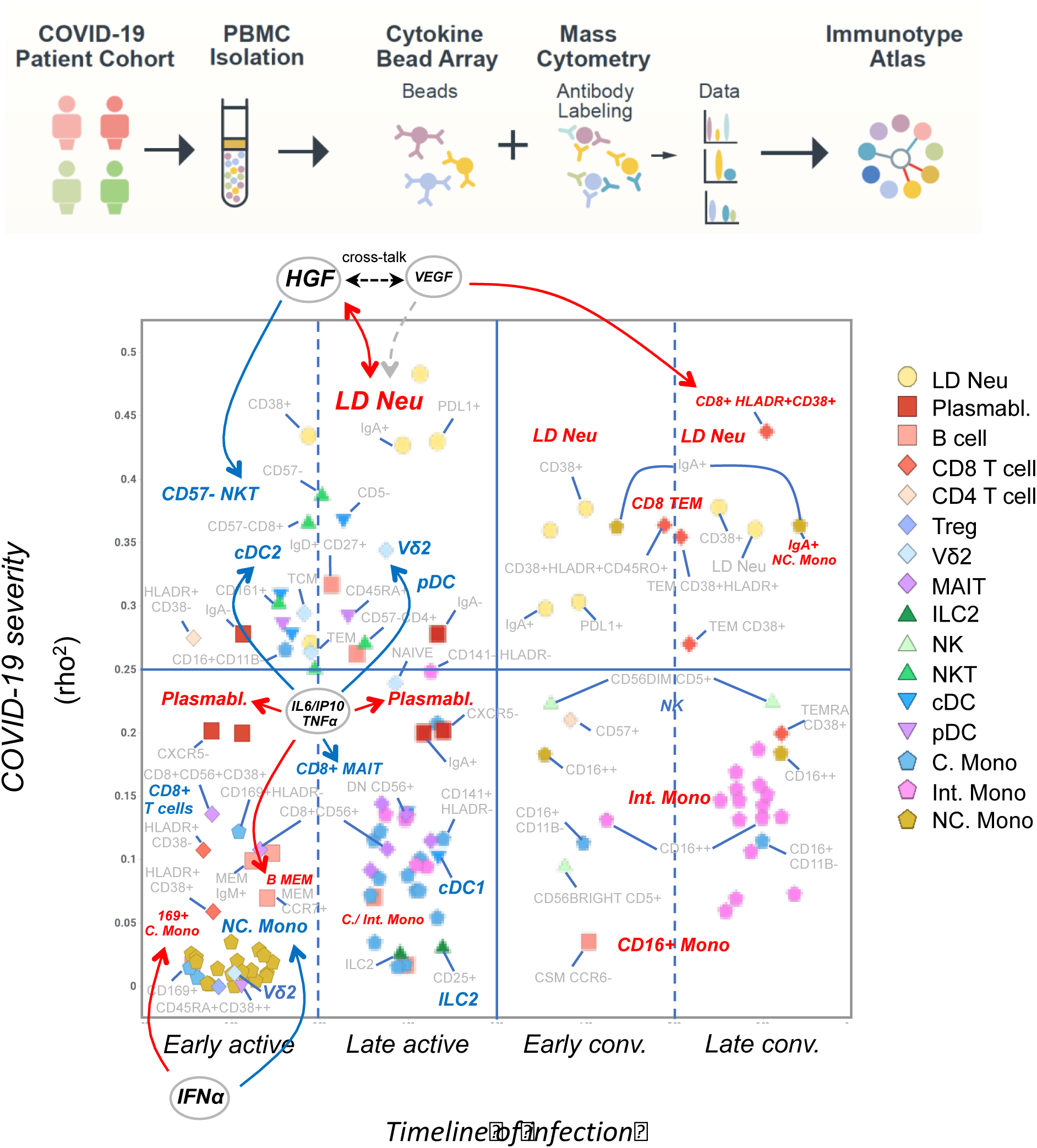
Timeline of infection and impact on COVID-19 disease severity. The figure is a summary of the timeline of infection versus the severity of COVID-19. The timeline is broken down into four phases (early active, late active, early conv. and late conv.), which were used to bin the extended set of 327 immune cells. The disease severity is expressed as the rho^2^ value from the correlation of their cell frequencies with the severity score. Shown are only cell subsets which have at least a 3-fold change in frequency in the respective phase (compare figures 7A and 7B) and/or a rho^2^ value > 0.25 for the disease correlation (compare figures 7C and 7D). The name of key cell subsets with increased blood frequencies are labeled in red, subsets with decreased frequency in blue. Cell subset/cytokine interactions are taken from figure 7.

**Table S1.** Demographic details of the study subjects. Population variables (sex and age at admission) and clinical variables (time from symptom onset to hospital admission, time from hospital admission to ICU entry, period of intubation, significant medical history and existing comorbidities) are shown. IQR, interquartile range; KW test, Kruskal-Wallis test; MW U test, Mann-Whitney U test; SD, standard deviation. * Includes the sole outlier who was intubated for 95 days before passing away from COVID-19. † Each instance of comorbidity was counted, even in cases where a patient had multiple comorbidities.

| Variables | Healthy Control<br>(N = 10) | COVID-19 |  |  |  |  |  |
| --- | --- | --- | --- | --- | --- | --- | --- |
|  |  | Total<br>(N = 77) | Statistical testing of total COVID-19 vs healthy controls (two-tailed p-value) | MILD (No O<sub>2</sub> Supplementation) - Groups I and VI (N = 31) | MODERATE (O<sub>2</sub> Supplementation) - Groups II and V (N = 20) | SEVERE (O<sub>2</sub> Supplementation + ICU) - Groups III and IV (N = 26) | Statistical testing of severity groups (two-tailed p-value, if applicable) |
| Sex - No. (%) |  |  |  |  |  |  |  |
| Male | 6 (60.0) | 60 (77.9) | 0.2128 (χ2 test) | 20 (64.5) | 17 (85.0) | 23 (88.5) | 0.0814 (χ2 test) |
| Female | 4 (40.0) | 17 (22.1) |  | 11 (35.5) | 3 (15.0) | 3 (11.5) |  |
| Age - Years |  |  |  |  |  |  |  |
| Mean ± SD. | 36.6 ± 9.2 | 50.4 ± 14.7 | 0.007 (MW U test) | 43.0 ± 13.0 | 52.9 ± 13.6 | 57.4 ± 13.9 | <0.0001 (KW test with Dunn's multiple comparison) |
| Median (IQR) | 34.0 (30.75-41.25) | 51.0 (39.5-62.0) |  | 44.0 (30.0-54.0) | 53.0 (42.5-62.5) | 60.5 (45.0-66.5) |  |
| Range | 25.0-56.0 | 24.0-82.0 |  | 24.0-65.0 | 28.0-80.0 | 29.0-82.0 |  |
| Time from Onset to Admission, Days |  |  |  |  |  |  |  |
| Mean ± SD. |  |  |  | 5.0 ± 8.1 | 5.4 ± 6.6 | 4.8 ± 3.4 | 0.528 (KW test with Dunn's multiple comparison) |
| Median (IQR) |  |  |  | 3.0 (1.0-5.0) | 4.0 (2.0-7.0) | 3.0 (2.0-9.0) |  |
| Range |  |  |  | 0.0-45.0 | 1.0-31.0 | 0.0-10.0 |  |

**Table S1.** Demographic details of the study subjects.
| Time from Admission to ICU, Days |  |  |  |  |  |  |  |
| --- | --- | --- | --- | --- | --- | --- | --- |
| Mean ± SD. |  |  |  |  |  | 2.4 ± 1.8 |  |
| Median (IQR) |  |  |  |  |  | 2.0 (1.0-4.0) |  |
| Range |  |  |  |  |  | 0.0-6.0 |  |
| Duration under Intubation, Days |  |  |  |  |  |  |  |
| Mean ± SD. |  |  |  |  |  | 7.9 ± 18.8 |  |
| Median (IQR) |  |  |  |  |  | 1.5 (0.0-7.75) |  |
| Range |  |  |  |  |  | 0.0-95.0* |  |
| Significant Medical History - No. (%) |  |  |  |  |  |  |  |
| Myocardial infarction |  | 12 (15.6) |  | 2 (6.5) | 5 (25.0) | 5 (19.2) | 0.1673 (χ2 test) |
| Comorbidity - No. (%)† |  |  |  |  |  |  |  |
| Hypertension |  | 27 (35.1) |  | 5 (16.1) | 9 (45.0) | 13 (50.0) | 0.8734 (χ2 test) |
| Hyperlipidemia/ dyslipidemia |  | 26 (33.8) |  | 3 (9.7) | 11 (55.0) | 12 (46.2) | 0.4603 (χ2 test) |
| Diabetes mellitus |  | 18 (23.4) |  | 1 (3.2) | 6 (30.0) | 11 (42.3) | 0.5487 (χ2 test) |

**Table S2.**
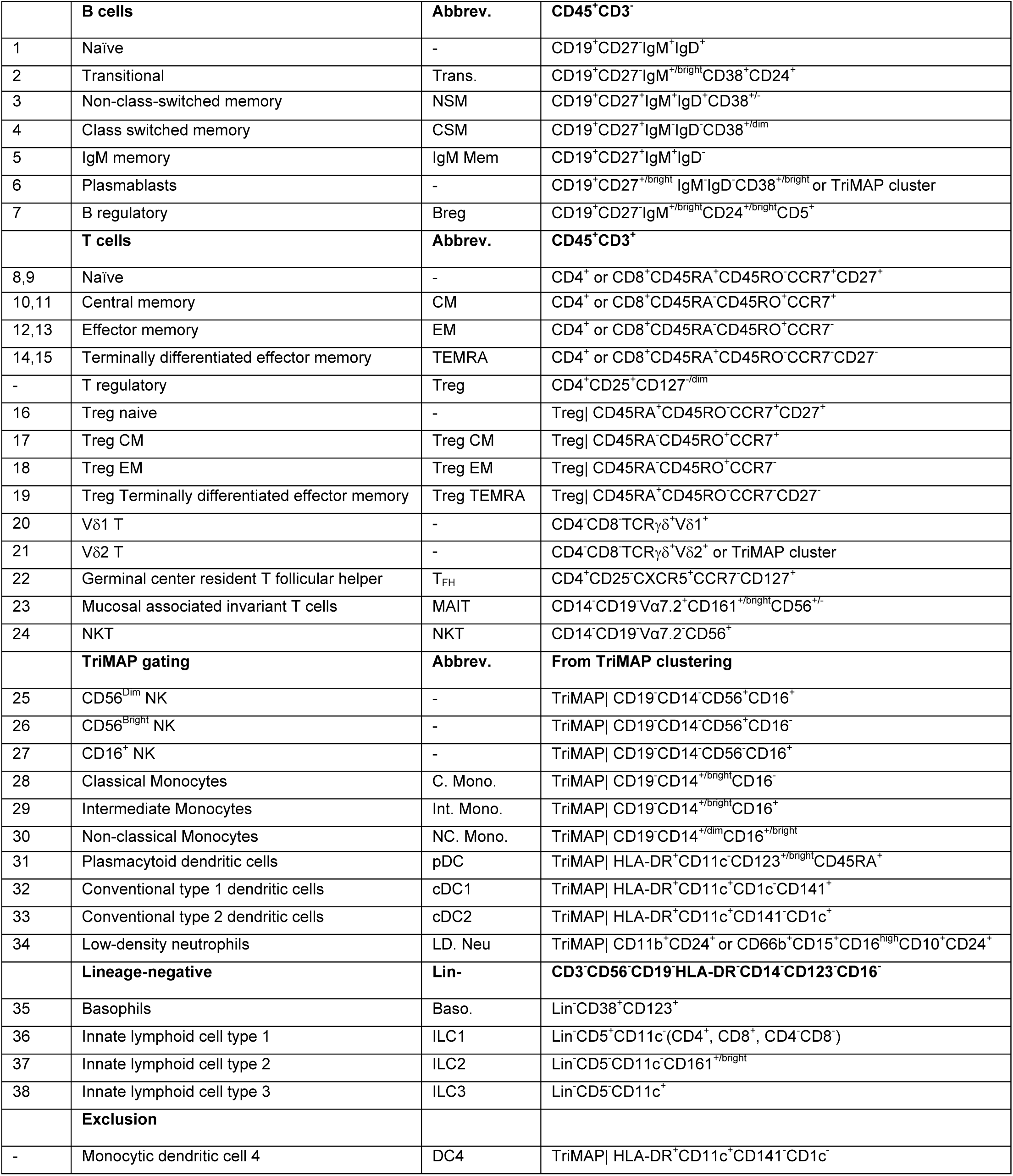
Gating strategy used for 38 basic immune subsets. For TriMAP gated subpopulations, the distinct clusters are calculated using TripMAP algorithm and confirmed based on their surface markers. (-) DC4 subset is excluded in analysis due to extreme low frequency.

**Table S3.** Definition of nodes used in timing and severity COVID-19 networks.

| Index | Alphabetical Order | Short name | Rank_num | Rank_name |
| --- | --- | --- | --- | --- |
| 1 | CytoF B CELLS | B CELLS | 45 | (45) B cells |
| 2 | CytoF B CELLS CCR7- | B CELLS CCR7- | 27 | (27) B CCR7- |
| 3 | CytoF B CELLS CCR7+ | B CELLS CCR7+ | 18 | (18) B CCR7+ |
| 4 | CytoF B CELLS CD38+ | B CELLS CD38+ | 3 | (3) B CD38+ |
| 5 | CytoF B CELLS CXCR5- | B CELLS CXCR5- | 8 | (8) B CXCR5- |
| 6 | CytoF B CELLS CXCR5+ | B CELLS CXCR5+ | 47 | (47) B CXCR5+ |
| 7 | CytoF B CELLS IgA+CD38++ | B CELLS IgA+CD38high | 7 | (7) B IgA+CD38++ |
| 8 | CytoF B CELLS IgD-CD27- | B CELLS IgD-CD27- | 16 | (16) B IgD-CD27- |
| 9 | CytoF B CELLS IgD+ | B CELLS IgD+ | 46 | (46) B IgD+ |
| 10 | CytoF B CELLS IgD+CD27+ | B CELLS IgD+CD27+ | 12 | (12) B IgD+CD27+ |
| 11 | CytoF B CELLS IgD+CD38+ | B CELLS IgD+CD38+ | 40 | (40) B IgD+CD38+ |
| 12 | CytoF B CELLS IgM+ | B CELLS IgM+ | 41 | (41) B IgM+ |
| 13 | CytoF B CELLS IgM+CD123+ | B CELLS IgM+CD123+ | 28 | (28) B IgM+CD123+ |
| 14 | CytoF B CELLS IgM+CD38++ | B CELLS IgM+CD38high | 10 | (10) B IgM+CD38++ |
| 15 | CytoF B CELLS IgA+ | B CELLS IgA+ | 32 | (32) B IgA+ |
| 16 | CytoF B CELLS MEM CCR7- | B CELLS MEM CCR7- | 24 | (24) B MEM CCR7- |
| 17 | CytoF B CELLS MEM CCR7+ | B CELLS MEM CCR7+ | 19 | (19) B MEM CCR7+ |
| 18 | CytoF B CELLS MEM CD38++ | B CELLS MEM CD38high | 2 | (2) B MEM CD38++ |
| 19 | CytoF B CELLS MEM CXCR5- | B CELLS MEM CXCR5- | 5 | (5) B MEM CXCR5- |
| 20 | CytoF B CELLS MEM CXCR5+ | B CELLS MEM CXCR5+ | 35 | (35) B MEM CXCR5+ |
| 21 | CytoF B CELLS MEM IgM+ | B CELLS MEM IgM+ | 36 | (36) B MEM IgM+ |
| 22 | CytoF B CELLS MEM IgM+CXCR5- | B CELLS MEM IgM+CXCR5- | 1 | (1) B MEM IgM+CXCR5- |
| 23 | CytoF B CELLS MEM IgM+CXCR5+ | B CELLS MEM IgM+CXCR5+ | 25 | (25) B MEM IgM+CXCR5+ |
| 24 | CytoF B CELLS NAIVE | B CELLS NAIVE | 42 | (42) B NAIVE |
| 25 | CytoF B CELLS NAIVE CCR7- | B CELLS NAIVE CCR7- | 37 | (37) B NAIVE CCR7- |
| 26 | CytoF B CELLS NAIVE CCR7+ | B CELLS NAIVE CCR7+ | 15 | (15) B NAIVE CCR7+ |
| 27 | CytoF B CELLS NAIVE CD123+ | B CELLS NAIVE CD123+ | 30 | (30) B NAIVE CD123+ |
| 28 | CytoF B CELLS NAIVE CD5- | B CELLS NAIVE CD5- | 17 | (17) B NAIVE CD5- |
| 29 | CytoF B CELLS NAIVE CD5+ | B CELLS NAIVE CD5+ | 22 | (22) B NAIVE CD5+ |
| 30 | CytoF B CELLS NAIVE CXCR5- | B CELLS NAIVE CXCR5- | 20 | (20) B NAIVE CXCR5- |
| 31 | CytoF B CELLS NAIVE IgM- | B CELLS NAIVE IgM- | 33 | (33) B NAIVE IgM- |
| 32 | CytoF B CELLS NAIVE IgM+ | B CELLS NAIVE IgM+ | 26 | (26) B NAIVE IgM+ |
| 33 | CytoF B MEM | B CELLS MEM | 34 | (34) B MEM |
| 34 | CytoF BASO | BASO | 1 | (1) BASO |
| 35 | CytoF BREG | BREG | 3 | (3) BREG |
| 36 | CytoF BREG CD123+ | BREG CD123+ | 1 | (1) BREG CD123+ |
| 37 | CytoF BREG CD38+ | BREG CD38+ | 2 | (2) BREG CD38+ |
| 38 | CytoF BREG PDL1 | BREG PDL1 | 4 | (4) BREG PDL1 |
| 39 | CytoF C. MONO | C. MONO | 17 | (17) C. Mono |
| 40 | CytoF C. Mono CD141- | C. Mono CD141- | 20 | (20) C. Mono CD141- |
| 41 | CyTOF C. Mono CD141-CD11B- | C. Mono CD141-CD11B- | 15 | (15) C. Mono CD141-CD11B- |
| 42 | CyTOF C. Mono CD141-CD11B+ | C. Mono CD141-CD11B+ | 18 | (18) C. Mono CD141-CD11B+ |
| 43 | CyTOF C. Mono CD141-HLADR- | C. Mono CD141-HLADR- | 7 | (7) C. Mono CD141-HLADR- |
| 44 | CyTOF C. Mono CD141-HLADR+ | C. Mono CD141-HLADR+ | 31 | (31) C. Mono CD141-HLADR+ |
| 45 | CyTOF C. Mono CD141+ | C. Mono CD141+ | 16 | (16) C. Mono CD141+ |
| 46 | CyTOF C. Mono CD141+CD11B- | C. Mono CD141+CD11B- | 13 | (13) C. Mono CD141+CD11B- |
| 47 | CyTOF C. Mono CD141+CD11B+ | C. Mono CD141+CD11B+ | 19 | (19) C. Mono CD141+CD11B+ |
| 48 | CyTOF C. Mono CD141+HLADR- | C. Mono CD141+HLADR- | 4 | (4) C. Mono CD141+HLADR- |
| 49 | CyTOF C. Mono CD141+HLADR+ | C. Mono CD141+HLADR+ | 27 | (27) C. Mono CD141+HLADR+ |
| 50 | CyTOF C. Mono CD16- | C. Mono CD16- | 11 | (11) C. Mono CD16- |
| 51 | CyTOF C. Mono CD16-CD11B- | C. Mono CD16-CD11B- | 22 | (22) C. Mono CD16-CD11B- |
| 52 | CyTOF C. Mono CD16-CD11B+ | C. Mono CD16-CD11B+ | 9 | (9) C. Mono CD16-CD11B+ |
| 53 | CyTOF C. Mono CD16-CD169- | C. Mono CD16-CD169- | 14 | (14) C. Mono CD16-CD169- |
| 54 | CyTOF C. Mono CD16-CD169+ | C. Mono CD16-CD169+ | 6 | (6) C. Mono CD16-CD169+ |
| 55 | CyTOF C. Mono CD16+ | C. Mono CD16+ | 24 | (24) C. Mono CD16+ |
| 56 | CyTOF C. Mono CD16+CD11B- | C. Mono CD16+CD11B- | 10 | (10) C. Mono CD16+CD11B- |
| 57 | CyTOF C. Mono CD16+CD11B+ | C. Mono CD16+CD11B+ | 26 | (26) C. Mono CD16+CD11B+ |
| 58 | CyTOF C. Mono CD16+CD169- | C. Mono CD16+CD169- | 23 | (23) C. Mono CD16+CD169- |
| 59 | CyTOF C. Mono CD16+CD169+ | C. Mono CD16+CD169+ | 25 | (25) C. Mono CD16+CD169+ |
| 60 | CyTOF C. Mono CD169-HLADR- | C. Mono CD169-HLADR- | 5 | (5) C. Mono CD169-HLADR- |
| 61 | CyTOF C. Mono CD169-HLADR+ | C. Mono CD169-HLADR+ | 28 | (28) C. Mono CD169-HLADR+ |
| 62 | CyTOF C. MONO CD169+ | C. MONO CD169+ | 3 | (3) C. Mono CD169+ |
| 63 | CyTOF C. Mono CD169+HLADR- | C. Mono CD169+HLADR- | 1 | (1) C. Mono CD169+HLADR- |
| 64 | CyTOF C. Mono CD169+HLADR+ | C. Mono CD169+HLADR+ | 29 | (29) C. Mono CD169+HLADR+ |
| 65 | CyTOF C. MONO CD38+ | C. MONO CD38+ | 21 | (21) C. Mono CD38+ |
| 66 | CyTOF C. MONO CD45RA- | C. MONO CD45RA- | 12 | (12) C. Mono CD45RA- |
| 67 | CyTOF C. MONO CD45RA+ | C. MONO CD45RA+ | 30 | (30) C. Mono CD45RA+ |
| 68 | CyTOF C. MONO CD56+ | C. MONO CD56+ | 8 | (8) C. Mono CD56+ |
| 69 | CyTOF C. MONO CD86+ | C. MONO CD86+ | 32 | (32) C. Mono CD86+ |
| 70 | CyTOF C. MONO IgA+ | C. MONO IgA+ | 2 | (2) C. Mono IgA+ |
| 71 | CyTOF CD141+HLADR-Cells | CD141+HLADR-Cells | 1 | (1) CD141+HLADR- Cells |
| 72 | CyTOF CD16+ NK | CD16+ NK | 1 | (1) CD16+ NK |
| 73 | CyTOF CD16+ NK CD38+ | CD16+ NK CD38+ | 2 | (2) CD16+ NK CD38+ |
| 74 | CyTOF CD4 CD11C+ | CD4 CD11C+ | 10 | (10) CD4 CD11C+ |
| 75 | CyTOF CD4 CD127+ | CD4 CD127+ | 18 | (18) CD4 CD127+ |
| 76 | CyTOF CD4 CD38+ | CD4 CD38+ | 24 | (24) CD4 CD38+ |
| 77 | CyTOF CD4 CD57+ | CD4 CD57+ | 33 | (33) CD4 CD57+ |
| 78 | CyTOF CD4 CXCR5- | CD4 CXCR5- | 20 | (20) CD4 CXCR5- |
| 79 | CyTOF CD4 CXCR5+ | CD4 CXCR5+ | 13 | (13) CD4 CXCR5+ |
| 80 | CyTOF CD4 HLADR-<br>CD38+CD45RO+ | CD4 HLADR- CD38+CD45RO+ | 34 | (34) CD4 HLADR-<br>CD38+CD45RO+ |
| 81 | CyTOF CD4 HLADR-CD38- | CD4 HLADR-CD38- | 12 | (12) CD4 HLADR-CD38- |
| 82 | CyTOF CD4 HLADR-CD38+ | CD4 HLADR-CD38+ | 21 | (21) CD4 HLADR-CD38+ |
| 83 | CyTOF CD4 HLADR+<br>CD38+CD45RO+ | CD4 HLADR+ CD38+CD45RO+ | 7 | (7) CD4 HLADR+<br>CD38+CD45RO+ |
| 84 | CyTOF CD4 HLADR+CD38- | CD4 HLADR+CD38- | 1 | (1) CD4 HLADR+CD38- |
| 85 | CyTOF CD4 HLADR+CD38+ | CD4 HLADR+CD38+ | 30 | (30) CD4 HLADR+CD38+ |
| 86 | CyTOF CD4 NAIVE | CD4 NAIVE | 15 | (15) CD4 NAIVE |
| 87 | CyTOF CD4 NAIVE CD127+ | CD4 NAIVE CD127+ | 16 | (16) CD4 NAIVE CD127+ |
| 88 | CyTOF CD4 NAIVE CD27+ | CD4 NAIVE CD27+ | 14 | (14) CD4 NAIVE CD27+ |
| 89 | CyTOF CD4 NAIVE CD38+ | CD4 NAIVE CD38+ | 4 | (4) CD4 NAIVE CD38+ |
| 90 | CyTOF CD4 T | CD4 T | 19 | (19) CD4 T |
| 91 | CyTOF CD4 TCM | CD4 TCM | 27 | (27) CD4 TCM |
| 92 | CyTOF CD4 TCM CD127+ | CD4 TCM CD127+ | 22 | (22) CD4 TCM CD127+ |
| 93 | CyTOF CD4 TCM CD27+ | CD4 TCM CD27+ | 25 | (25) CD4 TCM CD27+ |
| 94 | CyTOF CD4 TCM CD38+ | CD4 TCM CD38+ | 28 | (28) CD4 TCM CD38+ |
| 95 | CyTOF CD4 TCM CD38+HLADR+ | CD4 TCM CD38+HLADR+ | 3 | (3) CD4 TCM CD38+HLADR+ |
| 96 | CyTOF CD4 TEM | CD4 TEM | 29 | (29) CD4 TEM |
| 97 | CyTOF CD4 TEM CD127+ | CD4 TEM CD127+ | 26 | (26) CD4 TEM CD127+ |
| 98 | CyTOF CD4 TEM CD38+ | CD4 TEM CD38+ | 8 | (8) CD4 TEM CD38+ |
| 99 | CyTOF CD4 TEM CD38+HLADR+ | CD4 TEM CD38+HLADR+ | 5 | (5) CD4 TEM CD38+HLADR+ |
| 100 | CyTOF CD4 TEMRA | CD4 TEMRA | 17 | (17) CD4 TEMRA |
| 101 | CyTOF CD4 TEMRA CD27- | CD4 TEMRA CD27- | 2 | (2) CD4 TEMRA CD27- |
| 102 | CyTOF CD4 TEMRA CD38+ | CD4 TEMRA CD38+ | 31 | (31) CD4 TEMRA CD38+ |
| 103 | CyTOF CD4+ CD56- MAIT | CD4+ CD56- MAIT | 6 | (6) CD4+ CD56- MAIT |
| 104 | CyTOF CD4+ CD56- MAIT CD38+ | CD4+ CD56- MAIT CD38+ | 9 | (9) CD4+ CD56- MAIT CD38+ |
| 105 | CyTOF CD4+ CD56+ MAIT | CD4+ CD56+ MAIT | 23 | (23) CD4+ CD56+ MAIT |
| 106 | CyTOF CD4+ CD56+ MAIT CD38+ | CD4+ CD56+ MAIT CD38+ | 32 | (32) CD4+ CD56+ MAIT CD38+ |
| 107 | CyTOF CD4+CD25+ | CD4+CD25+ | 11 | (11) CD4+CD25+ |
| 108 | CyTOF CD56 BRIGHT NK | CD56 BRIGHT NK | 1 | (1) CD56 BRIGHT NK |
| 109 | CyTOF CD56 BRIGHT NK CD11B+ | CD56 BRIGHT NK CD11B+ | 4 | (4) CD56 BRIGHT NK CD11B+ |
| 110 | CyTOF CD56 BRIGHT NK CD11C+ | CD56 BRIGHT NK CD11C+ | 6 | (6) CD56 BRIGHT NK CD11C+ |
| 111 | CyTOF CD56 BRIGHT NK CD38+ | CD56 BRIGHT NK CD38+ | 5 | (5) CD56 BRIGHT NK CD38+ |
| 112 | CyTOF CD56 BRIGHT NK CD5+ | CD56 BRIGHT NK CD5+ | 2 | (2) CD56 BRIGHT NK CD5+ |
| 113 | CyTOF CD56 BRIGHT NK HLADR+ | CD56 BRIGHT NK HLADR+ | 3 | (3) CD56 BRIGHT NK HLADR+ |
| 114 | CyTOF CD56DIM NK | CD56DIM NK | 1 | (1) CD56DIM NK |
| 115 | CyTOF CD56DIM NK CD11B+ | CD56DIM NK CD11B+ | 2 | (2) CD56DIM NK CD11B+ |
| 116 | CyTOF CD56DIM NK CD11C+ | CD56DIM NK CD11C+ | 3 | (3) CD56DIM NK CD11C+ |
| 117 | CyTOF CD56DIM NK CD38+ | CD56DIM NK CD38+ | 4 | (4) CD56DIM NK CD38+ |
| 118 | CyTOF CD56DIM NK CD5+ | CD56DIM NK CD5+ | 5 | (5) CD56DIM NK CD5+ |
| 119 | CyTOF CD56DIM NK HLADR+ | CD56DIM NK HLADR+ | 6 | (6) CD56DIM NK HLADR+ |
| 120 | CyTOF CD8 CD11C+ | CD8 CD11C+ | 12 | (12) CD8 CD11C+ |
| 121 | CyTOF CD8 CD38+ | CD8 CD38+ | 23 | (23) CD8 CD38+ |
| 122 | CyTOF CD8 CXCR5- | CD8 CXCR5- | 16 | (16) CD8 CXCR5- |
| 123 | CyTOF CD8 CXCR5+ | CD8 CXCR5+ | 10 | (10) CD8 CXCR5+ |
| 124 | CyTOF CD8 HLADR- | CD8 HLADR- CD38+CD45RO+ | 24 | (24) CD8 HLADR- |

|  | CD38+CD45RO+ |  |  | CD38+CD45RO+ |
| --- | --- | --- | --- | --- |
| 125 | CyTOF CD8 HLADR-CD38- | CD8 HLADR-CD38- | 6 | (6) CD8 HLADR-CD38- |
| 126 | CyTOF CD8 HLADR-CD38+ | CD8 HLADR-CD38+ | 21 | (21) CD8 HLADR-CD38+ |
| 127 | CyTOF CD8 HLADR+<br>CD38+CD45RO+ | CD8 HLADR+ CD38+CD45RO+ | 26 | (26) CD8 HLADR+<br>CD38+CD45RO+ |
| 128 | CyTOF CD8 HLADR+CD38- | CD8 HLADR+CD38- | 1 | (1) CD8 HLADR+CD38- |
| 129 | CyTOF CD8 HLADR+CD38+ | CD8 HLADR+CD38+ | 28 | (28) CD8 HLADR+CD38+ |
| 130 | CyTOF CD8 NAIVE | CD8 NAIVE | 15 | (15) CD8 NAIVE |
| 131 | CyTOF CD8 NAIVE CD127+ | CD8 NAIVE CD127+ | 19 | (19) CD8 NAIVE CD127+ |
| 132 | CyTOF CD8 NAIVE CD27+ | CD8 NAIVE CD27+ | 17 | (17) CD8 NAIVE CD27+ |
| 133 | CyTOF CD8 NAIVE CD38+ | CD8 NAIVE CD38+ | 7 | (7) CD8 NAIVE CD38+ |
| 134 | CyTOF CD8 T CELL | CD8 T CELL | 13 | (13) CD8 T CELL |
| 135 | CyTOF CD8 TCM CD127+ | CD8 TCM CD127+ | 5 | (5) CD8 TCM CD127+ |
| 136 | CyTOF CD8 TCM CD27+ | CD8 TCM CD27+ | 14 | (14) CD8 TCM CD27+ |
| 137 | CyTOF CD8 TCM CD38+ | CD8 TCM CD38+ | 29 | (29) CD8 TCM CD38+ |
| 138 | CyTOF CD8 TCM CD38+HLADR+ | CD8 TCM CD38+HLADR+ | 27 | (27) CD8 TCM CD38+HLADR+ |
| 139 | CyTOF CD8 TEM | CD8 TEM | 20 | (20) CD8 TEM |
| 140 | CyTOF CD8 TEM CD127+ | CD8 TEM CD127+ | 3 | (3) CD8 TEM CD127+ |
| 141 | CyTOF CD8 TEM CD27+ | CD8 TEM CD27+ | 22 | (22) CD8 TEM CD27+ |
| 142 | CyTOF CD8 TEM CD38+ | CD8 TEM CD38+ | 30 | (30) CD8 TEM CD38+ |
| 143 | CyTOF CD8 TEM CD38+HLADR+ | CD8 TEM CD38+HLADR+ | 25 | (25) CD8 TEM CD38+HLADR+ |
| 144 | CyTOF CD8 TEMRA | CD8 TEMRA | 8 | (8) CD8 TEMRA |
| 145 | CyTOF CD8 TEMRA CD127+ | CD8 TEMRA CD127+ | 2 | (2) CD8 TEMRA CD127+ |
| 146 | CyTOF CD8 TEMRA CD27- | CD8 TEMRA CD27- | 4 | (4) CD8 TEMRA CD27- |
| 147 | CyTOF CD8 TEMRA CD38+ | CD8 TEMRA CD38+ | 11 | (11) CD8 TEMRA CD38+ |
| 148 | CyTOF CD8+ CD25+ | CD8+ CD25+ | 9 | (9) CD8+ CD25+ |
| 149 | CyTOF CD8+ CD56- MAIT | CD8+ CD56- MAIT | 2 | (2) MAIT CD8+ CD56- |
| 150 | CyTOF CD8+ CD56- MAIT CD38+ | CD8+ CD56- MAIT CD38+ | 1 | (1) MAIT CD8+ CD56- CD38+ |
| 151 | CyTOF CD8+ CD56+ MAIT | CD8+ CD56+ MAIT | 5 | (5) MAIT CD8+ CD56+ |
| 152 | CyTOF CD8+ CD56+ MAIT CD38+ | CD8+ CD56+ MAIT CD38+ | 10 | (10) MAIT CD8+ CD56+ CD38+ |
| 153 | CyTOF CD8+ TCM | CD8+ TCM | 18 | (18) CD8+ TCM |
| 154 | CyTOF cDC1 | cDC1 | 7 | (7) cDC1 |
| 155 | CyTOF cDC1 CD38+ | cDC1 CD38+ | 8 | (8) cDC1 CD38+ |
| 156 | CyTOF cDC2 | cDC2 | 2 | (2) cDC2 |
| 157 | CyTOF cDC2 CD11B+ | cDC2 CD11B+ | 6 | (6) cDC2 CD11B+ |
| 158 | CyTOF cDC2 CD38+ | cDC2 CD38+ | 3 | (3) cDC2 CD38+ |
| 159 | CyTOF cDC2 CD5- | cDC2 CD5- | 4 | (4) cDC2 CD5- |
| 160 | CyTOF cDC2 CD5+ | cDC2 CD5+ | 5 | (5) cDC2 CD5+ |
| 161 | CyTOF CSM | CSM | 43 | (43) B CSM |
| 162 | CyTOF CSM CCR6- | CSM CCR6- | 29 | (29) B CSM CCR6- |
| 163 | CyTOF CSM CCR6+ | CSM CCR6+ | 39 | (39) B CSM CCR6+ |
| 164 | CyTOF CSM CXCR5- | CSM CXCR5- | 31 | (31) B CSM CXCR5- |
| 165 | CyTOF CSM CXCR5+ | CSM CXCR5+ | 38 | (38) B CSM CXCR5+ |
| 166 | CyTOF DC | DC | 1 | (1) DC |
| 167 | CyTOF DC4 | DC4 | 9 | (9) cDC4 |
| 168 | CyTOF DC4 CD38+ | DC4 CD38+ | 10 | (10) cDC4 CD38+ |
| 169 | CyTOF DN CD56- MAIT | DN CD56- MAIT | 3 | (3) MAIT DN CD56- |
| 170 | CyTOF DN CD56- MAIT CD38+ | DN CD56- MAIT CD38+ | 6 | (6) MAIT DN CD56- CD38+ |
| 171 | CyTOF DN CD56+ MAIT | DN CD56+ MAIT | 4 | (4) MAIT DN CD56+ |
| 172 | CyTOF DN CD56+ MAIT CD38+ | DN CD56+ MAIT CD38+ | 11 | (11) MAIT DN CD56+ CD38+ |
| 173 | CyTOF ICOS+ TFH | ICOS+ TFH | 3 | (3) TFH ICOS+ |
| 174 | CyTOF ILC1 | ILC1 | 5 | (5) ILC1 |
| 175 | CyTOF ILC1 CD25- | ILC1 CD25- | 6 | (6) ILC1 CD25- |
| 176 | CyTOF ILC1 CD25+ | ILC1 CD25+ | 7 | (7) ILC1 CD25+ |
| 177 | CyTOF ILC1 CD38+ | ILC1 CD38+ | 8 | (8) ILC1 CD38+ |
| 178 | CyTOF ILC2 | ILC2 | 1 | (1) ILC2 |
| 179 | CyTOF ILC2 CD25- | ILC2 CD25- | 2 | (2) ILC2 CD25- |
| 180 | CyTOF ILC2 CD25+ | ILC2 CD25+ | 3 | (3) ILC2 CD25+ |
| 181 | CyTOF ILC2 CD38+ | ILC2 CD38+ | 4 | (4) ILC2 CD38+ |
| 182 | CyTOF ILC3 | ILC3 | 9 | (9) ILC3 |
| 183 | CyTOF ILC3 CD25- | ILC3 CD25- | 10 | (10) ILC3 CD25- |
| 184 | CyTOF ILC3 CD25+ | ILC3 CD25+ | 11 | (11) ILC3 CD25+ |
| 185 | CyTOF INT. MONO | INT. MONO | 14 | (14) INT. Mono |
| 186 | CyTOF Int. Mono CD141- | Int. Mono CD141- | 24 | (24) INT. Mono CD141- |
| 187 | CyTOF Int. Mono CD141-CD11B- | Int. Mono CD141-CD11B- | 18 | (18) INT. Mono CD141-CD11B- |
| 188 | CyTOF Int. Mono CD141-CD11B+ | Int. Mono CD141-CD11B+ | 17 | (17) INT. Mono CD141-CD11B+ |
| 189 | CyTOF Int. Mono CD141-HLADR- | Int. Mono CD141-HLADR- | 7 | (7) INT. Mono CD141-HLADR- |
| 190 | CyTOF Int. Mono CD141-HLADR+ | Int. Mono CD141-HLADR+ | 21 | (21) INT. Mono CD141-HLADR+ |
| 191 | CyTOF Int. Mono CD141+ | Int. Mono CD141+ | 11 | (11) INT. Mono CD141+ |
| 192 | CyTOF Int. Mono CD141+CD11B- | Int. Mono CD141+CD11B- | 10 | (10) INT. Mono CD141+CD11B- |
| 193 | CyTOF Int. Mono CD141+CD11B+ | Int. Mono CD141+CD11B+ | 16 | (16) INT. Mono CD141+CD11B+ |
| 194 | CyTOF Int. Mono CD141+HLADR- | Int. Mono CD141+HLADR- | 2 | (2) INT. Mono CD141+HLADR- |
| 195 | CyTOF Int. Mono CD141+HLADR+ | Int. Mono CD141+HLADR+ | 19 | (19) INT. Mono CD141+HLADR+ |
| 196 | CyTOF Int. Mono CD16+ | Int. Mono CD16+ | 8 | (8) INT. Mono CD16+ |
| 197 | CyTOF Int. Mono CD16++ | Int. Mono CD16++ | 22 | (22) INT. Mono CD16++ |
| 198 | CyTOF Int. Mono CD16+CD11B- | Int. Mono CD16+CD11B- | 13 | (13) INT. Mono CD16+CD11B- |
| 199 | CyTOF Int. Mono CD16+CD11B+ | Int. Mono CD16+CD11B+ | 15 | (15) INT. Mono CD16+CD11B+ |
| 200 | CyTOF Int. Mono CD16+CD169- | Int. Mono CD16+CD169- | 9 | (9) INT. Mono CD16+CD169- |
| 201 | CyTOF Int. Mono CD16+CD169+ | Int. Mono CD16+CD169+ | 25 | (25) INT. Mono CD16+CD169+ |
| 202 | CyTOF Int. Mono CD169-HLADR- | Int. Mono CD169-HLADR- | 4 | (4) INT. Mono CD169-HLADR- |
| 203 | CyTOF Int. Mono CD169-HLADR+ | Int. Mono CD169-HLADR+ | 12 | (12) INT. Mono CD169-HLADR+ |
| 204 | CyTOF INT. MONO CD169+ | INT. MONO CD169+ | 23 | (23) INT. Mono CD169+ |
| 205 | CyTOF Int. Mono CD169+HLADR- | Int. Mono CD169+HLADR- | 1 | (1) INT. Mono CD169+HLADR- |
| 206 | CyTOF Int. Mono CD169+HLADR+ | Int. Mono CD169+HLADR+ | 26 | (26) INT. Mono CD169+HLADR+ |
| 207 | CyTOF INT. MONO CD38+ | INT. MONO CD38+ | 20 | (20) INT. Mono CD38+ |
| 208 | CyTOF INT. MONO CD45RA- | INT. MONO CD45RA- | 6 | (6) INT. Mono CD45RA- |
| 209 | CyTOF INT. MONO CD45RA+ | INT. MONO CD45RA+ | 28 | (28) INT. Mono CD45RA+ |
| 210 | CyTOF INT. MONO CD56+ | INT. MONO CD56+ | 5 | (5) INT. Mono CD56+ |
| 211 | CyTOF INT. MONO CD86+ | INT. MONO CD86+ | 27 | (27) INT. Mono CD86+ |
| 212 | CyTOF INT. MONO IgA+ | INT. MONO IgA+ | 3 | (3) INT. Mono IgA+ |
| 213 | CyTOF LD Neu | LD Neu | 1 | (1) LD Neu |
| 214 | CyTOF LD Neu CD38+ | LD Neu CD38+ | 2 | (2) LD Neu CD38+ |
| 215 | CyTOF LD Neu CD5+ | LD Neu CD5+ | 3 | (3) LD Neu CD5+ |
| 216 | CyTOF LD Neu IgA+ | LD Neu IgA+ | 4 | (4) LD Neu IgA+ |
| 217 | CyTOF LD Neu PD-L1+ | LD Neu PD-L1+ | 5 | (5) LD Neu PD-L1+ |
| 218 | CyTOF MAIT | MAIT | 8 | (8) MAIT |
| 219 | CyTOF MAIT CD38+ | MAIT CD38+ | 13 | (13) MAIT CD38+ |
| 220 | CyTOF MAIT CD56- | MAIT CD56- | 9 | (9) MAIT CD56- |
| 221 | CyTOF MAIT CD56- CD38+ | MAIT CD56- CD38+ | 14 | (14) MAIT CD56- CD38+ |
| 222 | CyTOF MAIT CD56+ | MAIT CD56+ | 7 | (7) MAIT CD56+ |
| 223 | CyTOF MAIT CD56+ CD38+ | MAIT CD56+ CD38+ | 12 | (12) MAIT CD56+ CD38+ |
| 224 | CyTOF NC. MONO | NC. MONO | 16 | (16) NC. Mono |
| 225 | CyTOF NC. MONO CCR7- | NC. MONO CCR7- | 17 | (17) NC. Mono CCR7- |
| 226 | CyTOF NC. MONO CCR7+ | NC. MONO CCR7+ | 6 | (6) NC. Mono CCR7+ |
| 227 | CyTOF NC. MONO CD123+ | NC. MONO CD123+ | 13 | (13) NC. Mono CD123+ |
| 228 | CyTOF NC. Mono CD141- | NC. Mono CD141- | 20 | (20) NC. Mono CD141- |
| 229 | CyTOF NC. Mono CD141-CD11B- | NC. Mono CD141-CD11B- | 8 | (8) NC. Mono CD141-CD11B- |
| 230 | CyTOF NC. Mono CD141-CD11B+ | NC. Mono CD141-CD11B+ | 24 | (24) NC. Mono CD141-CD11B+ |
| 231 | CyTOF NC. Mono CD141-HLADR- | NC. Mono CD141-HLADR- | 2 | (2) NC. Mono CD141-HLADR- |
| 232 | CyTOF NC. Mono CD141-HLADR+ | NC. Mono CD141-HLADR+ | 21 | (21) NC. Mono CD141-HLADR+ |
| 233 | CyTOF NC. Mono CD141+ | NC. Mono CD141+ | 14 | (14) NC. Mono CD141+ |
| 234 | CyTOF NC. Mono CD141+CD11B- | NC. Mono CD141+CD11B- | 1 | (1) NC. Mono CD141+CD11B- |
| 235 | CyTOF NC. Mono CD141+CD11B+ | NC. Mono CD141+CD11B+ | 23 | (23) NC. Mono CD141+CD11B+ |
| 236 | CyTOF NC. Mono CD141+HLADR- | NC. Mono CD141+HLADR- | 7 | (7) NC. Mono CD141+HLADR- |
| 237 | CyTOF NC. Mono CD141+HLADR+ | NC. Mono CD141+HLADR+ | 15 | (15) NC. Mono CD141+HLADR+ |
| 238 | CyTOF NC. Mono CD16++ | NC. Mono CD16++ | 19 | (19) NC. Mono CD16++ |
| 239 | CyTOF NC. Mono CD16+CD11B- | NC. Mono CD16+CD11B- | 4 | (4) NC. Mono CD16+CD11B- |
| 240 | CyTOF NC. Mono CD16+CD11B+ | NC. Mono CD16+CD11B+ | 25 | (25) NC. Mono CD16+CD11B+ |
| 241 | CyTOF NC. Mono CD16+CD169- | NC. Mono CD16+CD169- | 12 | (12) NC. Mono CD16+CD169- |
| 242 | CyTOF NC. Mono CD16+CD169+ | NC. Mono CD16+CD169+ | 29 | (29) NC. Mono CD16+CD169+ |
| 243 | CyTOF NC. Mono CD169-HLADR- | NC. Mono CD169-HLADR- | 5 | (5) NC. Mono CD169-HLADR- |
| 244 | CyTOF NC. Mono CD169-HLADR+ | NC. Mono CD169-HLADR+ | 11 | (11) NC. Mono CD169-HLADR+ |
| 245 | CyTOF NC. Mono CD169+HLADR- | NC. Mono CD169+HLADR- | 18 | (18) NC. Mono CD169+HLADR- |
| 246 | CyTOF NC. Mono CD169+HLADR+ | NC. Mono CD169+HLADR+ | 27 | (27) NC. Mono CD169+HLADR+ |
| 247 | CyTOF NC. Mono CD16lo | NC. Mono CD16lo | 9 | (9) NC. Mono CD16lo |
| 248 | CyTOF NC. MONO CD38+ | NC. MONO CD38+ | 26 | (26) NC. Mono CD38+ |
| 249 | CyTOF NC. MONO CD45RO- | NC. MONO CD45RO- | 3 | (3) NC. Mono CD45RO- |
| 250 | CyTOF NC. MONO CD45RO+ | NC. MONO CD45RO+ | 30 | (30) NC. Mono CD45RO+ |
| 251 | CyTOF NC. MONO CD56+ | NC. MONO CD56+ | 22 | (22) NC. Mono CD56+ |
| 252 | CyTOF NC. MONO CD86+ | NC. MONO CD86+ | 10 | (10) NC. Mono CD86+ |
| 253 | CyTOF NC. MONO IgA+ | NC. MONO IgA+ | 28 | (28) NC. Mono IgA+ |
| 254 | CyTOF NK CD11B+ | NK CD11B+ | 1 | (1) NK CD11B+ |
| 255 | CyTOF NK CD11C+ | NK CD11C+ | 2 | (2) NK CD11C+ |
| 256 | CyTOF NKT | NKT | 1 | (1) NKT |
| 257 | CyTOF NKT CD161- | NKT CD161- | 2 | (2) NKT CD161- |
| 258 | CyTOF NKT CD161+ | NKT CD161+ | 3 | (3) NKT CD161+ |
| 259 | CyTOF NKT CD38+ | NKT CD38+ | 4 | (4) NKT CD38+ |
| 260 | CyTOF NKT CD57- | NKT CD57- | 5 | (5) NKT CD57- |
| 261 | CyTOF NKT CD57-CD4+ | NKT CD57-CD4+ | 6 | (6) NKT CD57-CD4+ |
| 262 | CyTOF NKT CD57-CD8+ | NKT CD57-CD8+ | 7 | (7) NKT CD57-CD8+ |
| 263 | CyTOF NKT CD57-DN | NKT CD57-DN | 8 | (8) NKT CD57-DN |
| 264 | CyTOF NKT CD57-DP | NKT CD57-DP | 9 | (9) NKT CD57-DP |
| 265 | CyTOF NKT CD57+ | NKT CD57+ | 10 | (10) NKT CD57+ |
| 266 | CyTOF NKT CD57+CD4+ | NKT CD57+CD4+ | 11 | (11) NKT CD57+CD4+ |
| 267 | CyTOF NKT CD57+CD8+ | NKT CD57+CD8+ | 12 | (12) NKT CD57+CD8+ |
| 268 | CyTOF NKT CD57+DN | NKT CD57+DN | 13 | (13) NKT CD57+DN |
| 269 | CyTOF NKT CD57+DP | NKT CD57+DP | 14 | (14) NKT CD57+DP |
| 270 | CyTOF NSM | NSM | 14 | (14) B NSM |
| 271 | CyTOF NSM CD27+CD38- | NSM CD27+CD38- | 6 | (6) B NSM CD27+CD38- |
| 272 | CyTOF NSM CD27+CD38+ | NSM CD27+CD38+ | 13 | (13) B NSM CD27+CD38+ |
| 273 | CyTOF NSM CD38+ | NSM CD38+ | 23 | (23) B NSM CD38+ |
| 274 | CyTOF PDC | PDC | 1 | (1) PDC |
| 275 | CyTOF PDC CD45RA+ | PDC CD45RA+ | 2 | (2) PDC CD45RA+ |
| 276 | CyTOF PDC CD45RA+CD38++ | PDC CD45RA+CD38high | 3 | (3) PDC CD45RA+CD38++ |
| 277 | CyTOF PLASMABLASTS | PLASMABLASTS | 3 | (3) PLASMABL. |
| 278 | CyTOF PLASMABLASTS CXCR5- | PLASMABLASTS CXCR5- | 2 | (2) PLASMABL. CXCR5- |
| 279 | CyTOF PLASMABLASTS CXCR5+ | PLASMABLASTS CXCR5+ | 5 | (5) PLASMABL. CXCR5+ |
| 280 | CyTOF PLASMABLASTS IgA- | PLASMABLASTS IgA- | 1 | (1) PLASMABL. IgA- |
| 281 | CyTOF PLASMABLASTS IgA+ | PLASMABLASTS IgA+ | 4 | (4) PLASMABL. IgA+ |
| 282 | CyTOF T CD38+ | T CD38+ | 6 | (6) T CD38+ |
| 283 | CyTOF T cells | T cells | 3 | (3) T cells |
| 284 | CyTOF T cells CCR7- | T cells CCR7- | 1 | (1) T CCR7- |
| 285 | CyTOF T cells CCR7+ | T cells CCR7+ | 5 | (5) T CCR7+ |
| 286 | CyTOF T CXCR5- | T CXCR5- | 4 | (4) T CXCR5- |
| 287 | CyTOF T CXCR5+ | T CXCR5+ | 2 | (2) T CXCR5+ |
| 288 | CyTOF TFH | TFH | 1 | (1) TFH |
| 289 | CyTOF TFH CD38+ | TFH CD38+ | 2 | (2) TFH CD38+ |
| 290 | CyTOF TRANS | TRANS | 11 | (11) B TRANS |
| 291 | CyTOF TRANS CCR6- | TRANS CCR6- | 44 | (44) B TRANS CCR6- |
| 292 | CyTOF TRANS CCR6+ | TRANS CCR6+ | 9 | (9) B TRANS CCR6+ |
| 293 | CyTOF TRANS CD5- | TRANS CD5- | 4 | (4) B TRANS CD5- |
| 294 | CyTOF TRANS CD5+ | TRANS CD5+ | 21 | (21) B TRANS CD5+ |
| 295 | CyTOF TREG | TREG | 3 | (3) TREG |
| 296 | CyTOF TREG CD11C+ | TREG CD11C+ | 1 | (1) TREG CD11C+ |
| 297 | CyTOF TREG CD38+ | TREG CD38+ | 11 | (11) TREG CD38+ |
| 298 | CyTOF TREG CD45RO-<br>ICOS+CD127+ | TREG CD45RO-ICOS+CD127+ | 6 | (6) TREG CD45RO-<br>ICOS+CD127+ |
| 299 | CyTOF TREG<br>CD45RO+ICOS+CD127+ | TREG CD45RO+ICOS+CD127+ | 9 | (9) TREG<br>CD45RO+ICOS+CD127+ |
| 300 | CyTOF TREG ICOS+ | TREG ICOS+ | 7 | (7) TREG ICOS+ |
| 301 | CyTOF TREG NAIVE | TREG NAIVE | 4 | (4) TREG NAIVE |
| 302 | CyTOF TREG NAIVE CD38+ | TREG NAIVE CD38+ | 5 | (5) TREG NAIVE CD38+ |
| 303 | CyTOF TREG TCM | TREG TCM | 2 | (2) TREG TCM |
| 304 | CyTOF TREG TCM CD38+ | TREG TCM CD38+ | 10 | (10) TREG TCM CD38+ |
| 305 | CyTOF TREG TEM | TREG TEM | 8 | (8) TREG TEM |
| 306 | CyTOF TREG TEM CD38+ | TREG TEM CD38+ | 13 | (13) TREG TEM CD38+ |
| 307 | CyTOF TREG TEMRA | TREG TEMRA | 12 | (12) TREG TEMRA |
| 308 | CyTOF VD1 | VD1 | 11 | (11) VD1 |
| 309 | CyTOF VD1 CD38+ | VD1 CD38+ | 12 | (12) VD1 CD38+ |
| 310 | CyTOF VD1 NAIVE | VD1 NAIVE | 13 | (13) VD1 NAIVE |
| 311 | CyTOF VD1 NAIVE CD38+ | VD1 NAIVE CD38+ | 14 | (14) VD1 NAIVE CD38+ |
| 312 | CyTOF VD1 TCM | VD1 TCM | 15 | (15) VD1 TCM |
| 313 | CyTOF VD1 TCM CD38+ | VD1 TCM CD38+ | 16 | (16) VD1 TCM CD38+ |
| 314 | CyTOF VD1 TEM | VD1 TEM | 17 | (17) VD1 TEM |
| 315 | CyTOF VD1 TEM CD38+ | VD1 TEM CD38+ | 18 | (18) VD1 TEM CD38+ |
| 316 | CyTOF VD1 TEMRA | VD1 TEMRA | 19 | (19) VD1 TEMRA |
| 317 | CyTOF VD1 TEMRA CD38+ | VD1 TEMRA CD38+ | 20 | (20) VD1 TEMRA CD38+ |
| 318 | CyTOF VD2 | VD2 | 1 | (1) VD2 |
| 319 | CyTOF VD2 CD38++ | VD2 CD38high | 2 | (2) VD2 CD38++ |
| 320 | CyTOF VD2 NAIVE | VD2 NAIVE | 3 | (3) VD2 NAIVE |
| 321 | CyTOF VD2 NAIVE CD38+ | VD2 NAIVE CD38+ | 4 | (4) VD2 NAIVE CD38+ |
| 322 | CyTOF VD2 TCM | VD2 TCM | 5 | (5) VD2 TCM |
| 323 | CyTOF VD2 TCM CD38+ | VD2 TCM CD38+ | 6 | (6) VD2 TCM CD38+ |
| 324 | CyTOF VD2 TEM | VD2 TEM | 7 | (7) VD2 TEM |
| 325 | CyTOF VD2 TEM CD38+ | VD2 TEM CD38+ | 8 | (8) VD2 TEM CD38+ |
| 326 | CyTOF VD2 TEMRA | VD2 TEMRA | 9 | (9) VD2 TEMRA |
| 327 | CyTOF VD2 TEMRA CD38+ | VD2 TEMRA CD38+ | 10 | (10) VD2 TEMRA CD38+ |

